# Prefrontal co-expression of schizophrenia risk genes is associated with treatment response in patients

**DOI:** 10.1101/323428

**Authors:** Giulio Pergola, Pasquale Di Carlo, Andrew E. Jaffe, Marco Papalino, Qiang Chen, Thomas M. Hyde, Joel E. Kleinman, Joo Heon Shin, Antonio Rampino, Giuseppe Blasi, Daniel R. Weinberger, Alessandro Bertolino

**Affiliations:** Group of Psychiatric Neuroscience, Department of Basic Medical Sciences, Neuroscience and Sense Organs, University of Bari Aldo Moro, Bari, Italy; Lieber Institute for Brain Development, Johns Hopkins Medical Campus, Baltimore, Maryland, USA; Department of Mental Health, Johns Hopkins Bloomberg School of Public Health, Baltimore, Maryland, USA; Department of Biostatistics, Johns Hopkins Bloomberg School of Public Health, Baltimore, Maryland, USA; Center for Computational Biology, Johns Hopkins University, Baltimore, Maryland, USA; Department of Neuroscience, Johns Hopkins School of Medicine, Baltimore, Maryland, USA; Department of Neurology, Johns Hopkins School of Medicine, Baltimore; Department of Psychiatry and Behavioral Sciences, Johns Hopkins School of Medicine, Baltimore, Maryland, USA; McKusick-Nathans Institute of Genetic Medicine, Johns Hopkins School of Medicine, Baltimore, Maryland, USA; Azienda Ospedaliero-Universitaria Consorziale Policlinico, Bari, Italy

**Keywords:** Gene co-expression networks, dorsolateral prefrontal cortex, olanzapine, RNA sequencing, schizophrenia

## Abstract

Gene co-expression networks are relevant to functional and clinical translation of schizophrenia (SCZ) risk genes. We hypothesized that SCZ risk genes may converge into coexpression pathways which may be associated with gene regulation mechanisms and with response to treatment in patients with SCZ. We identified gene co-expression networks in two prefrontal cortex *post-mortem* RNA sequencing datasets (total N=688) and replicated them in four more datasets (total N=227). We identified and replicated (all p-values<.001) a single module enriched for SCZ risk loci (13 risk genes in 10 loci). *In silico* screening of potential regulators of the SCZ risk module via bioinformatic analyses identified two transcription factors and three miRNAs associated with the risk module. To translate *post-mortem* information into clinical phenotypes, we identified polymorphisms predicting co-expression and combined them to obtain an index approximating module co-expression (Polygenic Co-expression Index: PCI). The PCI-co-expression association was successfully replicated in two independent brain transcriptome datasets (total N=131; all p-values<.05). Finally, we tested the association between the PCI and short-term treatment response in two independent samples of patients with SCZ treated with olanzapine (total N=167). The PCI was associated with treatment response in the positive symptom domain in both clinical cohorts (all p-values<.05).

In summary, our findings in a large sample of human *post-mortem* prefrontal cortex show that coexpression of a set of genes enriched for schizophrenia risk genes is relevant to treatment response. This co-expression pathway may be co-regulated by transcription factors and miRNA associated with it.

## Introduction

Schizophrenia (SCZ) risk is highly related to genetic factors, and specific risk loci have recently been identified by the Psychiatric Genomics Consortium (PGC) (*1*). The discovery that at least 108 genetic loci are associated with the disease suggests that multiple biological processes may be involved in SCZ, perhaps converging into one or few common pathways (high coherence), or distributed across many pathways of genetic risk (low coherence) (*2*). The question of genetic risk coherence is an important issue in SCZ research because the functional and clinical translation of PGC SCZ risk variants remains modest when they are considered on their own or additively cumulated. For example, currently available cumulative scores do not explain a large fraction of the variance in treatment response and treatment resistance (*3*, *4*).

The challenge of translating genetic risk into common pathways associated with clinical predictions is compounded by the fact that we know the risk ***loci***, but in only a minority of cases do we know which ***genes*** within them are causally implicated in the disorder. PGC risk loci include many genes and are proximal to many more, such that risk variants in the loci may theoretically impact hundreds of genes (*5*); additionally, the effect of genetic variants in the PGC loci is not necessarily restricted to proximal genes (*6*). Understanding the relationship between risk variants and genes involved in the disorder may require identification of ***common pathways and biological processes involving genes located in multiple loci*** - rather than considering only the genetic variants associated with GWAS hits. In turn, discovering biological pathways that bring together multiple SCZ risk loci will contribute to identify molecular elements, such as transcription factors and miRNA, that may represent ***nodes of risk convergence*** by regulating diverse gene functions.

A basic principle of biology is that the expression of individual genes is often coordinated by regulatory molecules resulting in the co-expression of gene networks (*7*). Therefore, gene coexpression is a biological process possibly relevant to the convergence of SCZ risk into common pathways that are associated with clinical translation of PGC loci. At least some of the PGC risk variants control gene expression (*8, 9*). Recent evidence suggests that genes in the PGC SCZ loci co-segregate into co-expression pathways (*8*) and genetic variation in such pathways is relevant to SCZ phenotypes (*10*).

We hypothesized that ***genes located in PGC SCZ risk loci may converge into co-expression pathways*** which, in turn, may reveal molecular elements potentially contributing to orchestrate genetic risk into common biological pathways and ultimately clinical outcome. The translational relevance of such co-expression pathways to SCZ can be validated in terms of their association with clinical phenotypes in patients, including treatment outcome. However, gene set clustering in co-expression networks is variable and methodologically complex (*11*), and thus requires transcriptome-wide replication to be considered reliable. Fromer and coworkers (*8*) previously used RNA sequencing in *post-mortem* prefrontal cortex to identify gene expression patterns potentially relevant to SCZ risk. Here, we used RNA sequencing data from the two largest collections of *post-mortem* prefrontal cortex currently available: the Lieber Institute for Brain Development repository (LIBD) (*12*) and the CommonMind Consortium collection (CMC) (*8*). We identified gene co-expression networks by means of Weighted Gene Co-expression Network Analysis (WGCNA) (*13*). After assessing network preservation of the LIBD co-expression network in CMC, we focused on one module showing overrepresentation of genes located in the PGC SCZ loci. We aimed to identify potential genetic regulators of the loci and to assess clinical translation. In order to translate *post-mortem* data mining into clinical phenotypes, we used common genetic variation, i.e., we identified co-expression quantitative trait loci (co-eQTLs) (*10*) and combined them to obtain a numeric index approximating network co-expression (*14*). Genetic variation in this gene set was associated with short-term treatment response to olanzapine in terms of positive symptoms in the largest double-blind clinical trial openly available to date with genome-wide genotyping (CATIE; N = 121) (*15*). We replicated the clinical results in an independent dataset of 46 patients with SCZ treated with olanzapine in Bari, Italy (*16*). The current work complements further reports on partially overlapping datasets which focused on network approaches to identify potential novel drug targets (*17*).

## Results

### Co-expression Network of human prefrontal cortex

We selected frontal cortex samples from 343 LIBD subjects and 345 CMC subjects. The sample was filtered based on RNA Integrity Number (≥7.0), age range (17-86 years), ethnicity (African-American and Caucasians), and diagnosis (LIBD: patients with SCZ = 143, healthy controls [HCs] = 200; CMC: patients with SCZ = 166; HCs = 179; demographics in Table 1). Transcripts available in both datasets with Reads Per Kilobase per Million (RPKM) > .1 mapped to 20,993 genes. After preprocessing (*11*), we computed WGCNA, separately for patients with SCZ and HCs within each of the two datasets, and derived network preservation statistics (Fig. S1) (*18*, *19*). We found that all co-expression modules showed moderate to strong preservation between patients with SCZ and HCs both in the LIBD and in CMC datasets (all Z-summary scores ≥ 2, Fig. S1). Since all modules were thus relatively preserved between patients with SCZ and HCs across both datasets at the selected threshold (results in Supplementary Materials, SM, and Fig. S1), we pooled data from patients and controls and identified one network for the LIBD and one for the CMC datasets. The results reported in the manuscript refer to this WGCNA with pooled patients with SCZ and HCs. In this WGCNA, we selected the LIBD network as the reference and tested its preservation in the CMC network, which was successful (Fig. 1A-B; additional details in the SM; Table S1 and S2; Fig. S2). The whole network identified in the LIBD dataset with gene modules and connectivity statistics is available in Data file S1.

**Table 1.** Demographics.

|  | LIBD | CMC | Stats LIBD vs. CMC |  | BRAINEAC | LIBD<br>Developmental<br>Ages | GTEx:<br>Brain Cortex | GTEx:<br>Brain BA9 | CATIE | UNIBA |
| --- | --- | --- | --- | --- | --- | --- | --- | --- | --- | --- |
| **Sample size** | 343 | 345 |  |  | 38 | 93 | 96 | 92 | 121 | 46 |
| **Female (male)**<br>**[ratio]** | 105 (238)<br>[0.44] | 114 (231)<br>[0.49] | $\chi^2 = 0.4$ | $p = 0.55$ | 9 (29) [0.31] | 33 (60) [0.55] | # | # | 27 (94)<br>[0.28] | 9 (37)<br>[0.24] |
| **Age mean ± s.d.**<br>**(years)** | 45.2 ± 14.8 | 60.8 ± 17.4 | $t = -12.7$ | $p < 2.2 \times 10^{-16}$ | 56.6 ± 19.1 | 3.5 ± 6.0 | # | # | 41.5 ± 10.9 | 27.7 ± 6.8 |
| **Age range (years)** | 17-85 | 17-86 |  |  | 20-89 | 0-16 | # | # | 19-65 | 16-42 |
| **Diagnosis SCZ<sup>a</sup>**<br>**(HC<sup>b</sup>)** | 143 (200) | 166 (179) | $\chi^2 = 2.6$ | $p = 0.11$ | 0 (38) | 0 (93) | # | # | 121 (0) | 46 (0) |
| **Ethnicity CAUC<sup>c</sup>**<br>**(AA<sup>d</sup>)** | 166 (177) | 283 (62) | $\chi^2 = 84.3$ | $p < 2.2 \times 10^{-16}$ | 38 (0) | 40 (53) | # | # | 77 (44) | 46 (0) |

<sup>a</sup> patients with schizophrenia; <sup>b</sup> healthy controls; <sup>c</sup> Caucasian; <sup>d</sup> African-American; # data unavailable as per ref. 49.

### Prioritization of modules relevant for SCZ

We prioritized modules in terms of their relevance for SCZ. To assess their relevance to diagnosis, we tested whether any module eigengenes (the first principal component of gene expression in the modules, abbreviated as *ME* in the following) were associated with diagnosis. Six *ME* were significantly different between patients with SCZ and HCs in the LIBD network (Bonferroni-corrected p-value < .05, Table S3), suggesting potentially different co-expression. However, none of these associations were replicated in CMC (all uncorrected p-values > .05, Table S3). Having found no replicable co-expression signature of diagnosis, we asked whether any of the modules included more risk genes for SCZ than expected by chance. This way we ***directly tested the hypothesis that risk converges into co-expression pathways***. To detect modules in which PGC SCZ risk genes (n = 310; gene list in the Table S4) were overrepresented, we computed hypergeometric tests and corrected the results for multiple comparisons (Bonferroni-corrected p-value < .05). We found that the *Darkgreen* module in the LIBD network was the only module significantly enriched for genes in the PGC SCZ loci (10 loci, 13 genes, p-value = 3.1×10^−5^, see Table 2, see Table S5 for the full list of *Darkgreen* genes). Notably, the enrichment remained significant when including both protein-coding and nonprotein-coding genes located in the PGC loci (p-value = 5.7×10^-4^), further suggesting that *Darkgreen* co-expressed genes co-localized with genetic risk for SCZ. Next, we asked whether this enrichment was affected by genetic spatial proximity. Our hypothesis was that the overrepresentation of SCZ risk genes should remain significant when expanding the boundaries of the loci within a genomic distance compatible with an influence of sequence elements on gene expression (*20*, *21*). The enrichment survived permutation-based empirical p-value < .001 when loci were expanded up to 450 kbp (Fig. 1C; see also Fig. S3, which includes protein-coding and non-protein-coding genes), indicating that many genes in the same loci were co-expressed in *Darkgreen.* Additionally, gene set ‘competitive’ enrichment analysis with the software MAGMA (*22*) demonstrated that variants falling within *Darkgreen* were associated with greater SCZ risk compared to the remaining sets (we excluded the *Grey* module of non-clustered genes; p-value = .036, see Methods and Table S1 for details). Hence, converging evidence from the gene list and the localization of genetic variants suggested that genetic risk for SCZ converged into *Darkgreen*. Moreover, *ME Darkgreen* was not associated with possible biological confounders such as smoking habit, nor with antipsychotic or antidepressant medications in SCZ patients (we used a binary classification of whether or not patients used the substances; uncorrected p-value > .1; Data file S2 reports uncorrected p-value for all modules in the LIBD network).

### Functional significance of Darkgreen module

*Darkgreen* included 225 genes, of which 157 were protein coding (Table S5). We investigated the functional significance of *Darkgreen* by means of gene ontology analyses. *Darkgreen* was functionally enriched for gene products involved in homophilic cell adhesion via plasma membrane (Amigo2, GO:0007156, 9 genes, fold-enrichment = 7.92, Bonferroni-corrected p-value = .022). Specific Expression Analysis [http://genetics.wustl.edu/jdlab/csea-tool-2/] (*23*) revealed that *Darkgreen* was enriched for genes preferentially expressed in the cortex during young adulthood (*24*) (Fig. S4). Therefore, we asked whether *Darkgreen* genes were also co-expressed during neurodevelopment, given the importance of developmental ages for SCZ liability (*25*). WGCNA on a sample of 93 LIBD subjects from fetuses to 16 year old individuals (hereinafter, LIBD developmental series) nonoverlapping with the sample used in the main analysis revealed higher than chance topological preservation (empirical p-value < .001 (*26*); Table S6), showing that *Darkgreen* gene-gene relationships were significant also in independent subjects during developmental life stages. We also explored further datasets to assess the robustness of the gene-gene relationships detected in this module. *Darkgreen* was among 12 modules preserved in all of the three additional frontal cortex microarray and RNA sequencing datasets we analyzed, showing that the gene-gene associations we identified were robust (Fig. 1D; empirical p-value < .001 (*26*); Table S6). Fig.1E represents the hub genes of *Darkgreen* and their relationship with PGC SCZ hits included in the module.

### Genetic regulation potentially implicated in Darkgreen module co-expression

We hypothesized that co-expressed genes may be co-regulated by elements such as transcription factors (TFs) and miRNA. We tested this hypothesis by investigating transcription factors targeting *Darkgreen* genes. Using the software Pscan (http://159.149.160.88/pscan/)(*27*) we identified two TFs (NRF1, KLF14) whose binding motif was overrepresented in the promoter regions of our co-expressed genes (Bonferroni-corrected p-value < .05). Interestingly, seven out of 13 SCZ risk genes showed an association with NRF1 (*GIGYF2*, *NDUFA6*, *SCAF1*, *CACNA1C*, *IGSF9B*, *TMX2*, *ANKRD44*); also KLF14 had seven PGC risk gene targets (*SCAF1*, *ANKRD44*, *GIGYF2*, *TMX2*, *CACNA1C*, *IGSF9B*, *AKT3*). However, the identified TF were related with several other modules (corrected p-value < .05; NRF1 to 24 modules; KLF14 to 18 modules; Fig. S5), hindering conclusions about their specificity. It is also possible that some TFs may exert their effects on multiple modules because of tissue expression specificity or biological coherence of the identified modules.

Micro-RNAs (miRNAs) are also regulators of gene co-expression (*28*). Hauberg and coworkers (*29*) have shown that the targetome of 10 miRNAs is enriched for SCZ risk variants. Here, we assessed the overrepresentation of the targetome of each of these miRNAs in *Darkgreen*. We found that the targets of three SCZ-related miRNA (miR-101, miR-374, miR-28) were overrepresented in *Darkgreen* (Bonferroni-corrected p-value < .05; see Table S7 for further details), suggesting that these miRNAs may plausibly promote the correlated expression of *Darkgreen* genes. Both miR-374 and miR-28 targets shared the same seven SCZ risk genes (*AKT3*, *ANKRD44*, *CACNA1C*, *PCDHA3*, *PCDHA4*, *PCDHA5*, *PCDHA6*), while among miR-101 targets we found two risk genes (*ANKRD44* and *AKT3*). To assess the specificity of these findings, we computed the enrichment for miRNA targetomes in all other modules and reported uncorrected p-values in Data file S3. The identified miRNAs overlapped with only few modules (miR-101/miR-374/miR-28 = 8/6/10 modules, corrected p-value < .05, Fig. S6), suggesting some degree of specificity. Overall, these results are consistent with the idea that genetic risk convergence in *Darkgreen* may be mediated by TFs and miRNAs.

### Overlap with genes regulated by antipsychotics

We defined our network based on data from patients with SCZ and HCs. Since SCZ patients are usually treated with antipsychotics, it can be hypothesized that drugs contributed to the aggregation of genes into modules. In a recent study, Kim and coworkers (*30*) identified genes differentially expressed in the striatum and in the whole brain of mice exposed to haloperidol vs. not exposed mice. We computed for all modules the enrichment for the differentially expressed genes and found a single module (*Brown*) enriched (11 genes, 16% of total differentially expressed genes, Bonferroni-corrected p-value = .00447, Data file S4). Specifically, *Brown* was enriched for down-regulated genes (9 genes, 26.5% of down-regulated genes, Bonferroni-corrected p-value = .00411; Fig. S7). *Darkgreen* did not show any significant overlap with haloperidol target genes, suggesting that its relevance for SCZ risk genes was not a by-product of medication, at least to the extent that haloperidol is a representative antipsychotic.

**Fig. 1.**
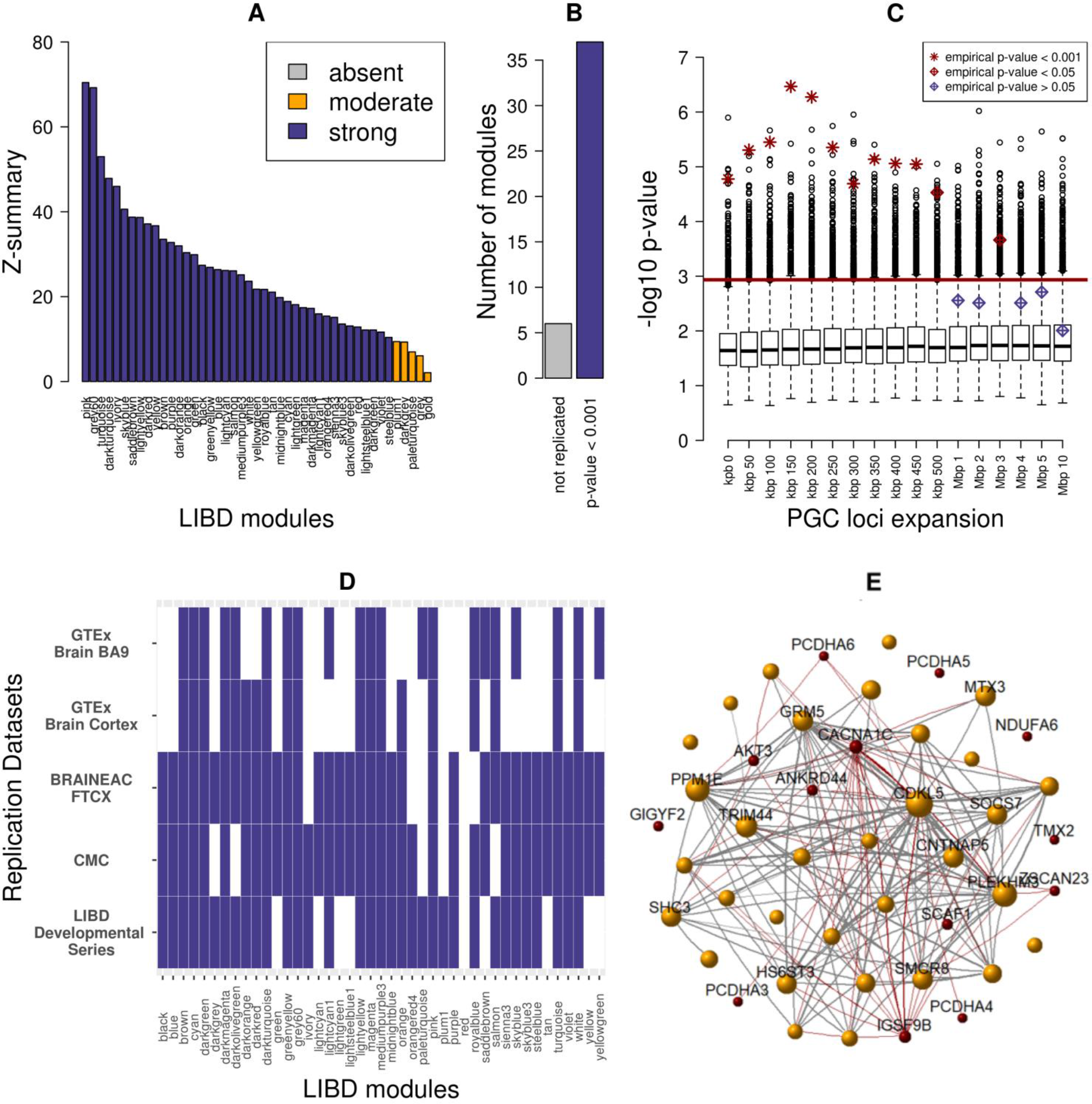
Co-expression Network. (**A**) Preservation of the LIBD network in the CMC dataset (Langfelder method). The LIBD modules are shown on the x-axis ranked by Z-summary preservation score (y-axis). Z ≥ 10 denotes strong preservation, 2 ≤ Z < 10 moderate, and Z < 2 absent (*18*). (**B**) Replication of the LIBD modules topology in the CMC dataset (Johnson method). Bars indicate the 44 number of replicated modules at empirical p-value < .001 vs. not replicated modules (10,000 permutations). (**C**) *Darkgreen* module enrichment for schizophrenia risk genes. Enrichment significance is shown over increasing expansion of schizophrenia risk loci boundaries. The x-axis reports the size of expansion in kilo-base pairs (kbp). The y-axis indicates the −log_10_ p-value of the hypergeometric test for overrepresentation of schizophrenia risk loci in *Darkgreen*. Boxplots show the null distribution of the lowest enrichment p-value over all network modules obtained after network labels permutation (n=10,000). The red horizontal line shows the Bonferroni threshold selected (number of modules = 43, α = .0012). Stars and diamonds denote *Darkgreen* exact enrichment p-value. (**D**) Replication of the LIBD modules in several different datasets (Johnson method). Slate-blue fields denote modules (x-axis) replicated at empirical p-value < .001 (over 10,000 permutations). (**E**) *Darkgreen* graph. The nodes of the graphs (spheres) are genes and schizophrenia risk genes are colored in dark red. Gold spheres represent a selection of the most connected genes in the module (scaled intra-modular connectivity ≥ 0.3) and have a diameter proportional to intra-modular connectivity. i.e., larger spheres denote genes harboring more connections within *Darkgreen*. Lines denote gene-gene relationships and their width is proportional to connection strength.

**Table 2.** PGC loci and genes overlapping with the module *Darkgreen*.

| Ensembl gene ID | HGNC<sup>a</sup><br>Symbol | Gene name | PGC loci<br>rank | PGC loci<br>index SNP | PGC loci<br>position (hg19<sup>b</sup>) | PGC<br>index SNP<br>p-value |
| --- | --- | --- | --- | --- | --- | --- |
| ENSG00000187987 | ZSCAN23 | zinc finger and SCAN domain containing 23 | 1 | rs115329265 | chr6:28303247-28712247 | 3.48×10<sup>-31</sup> |
| ENSG00000151067 | CACNA1C | calcium channel, voltage-dependent, L type, alpha 1C subunit | 4 | rs2007044, rs2239063 | chr12:2321860-2523731 | 3.22×10<sup>-18</sup> |
| ENSG00000204120 | GIGYF2 | GRB10 interacting GYF protein 2 | 22 | rs6704768 | chr2:233559301-233753501 | 2.32×10<sup>-12</sup> |
| ENSG00000065413 | ANKRD44 | ankyrin repeat domain 44 | 31 | rs6434928 | chr2:198148577-198835577 | 2.06×10<sup>-11</sup> |
| ENSG00000080854 | IGSF9B | immunoglobulin superfamily, member 9B | 36 | rs75059851 | chr11:133808069-133852969 | 3.87×10<sup>-11</sup> |
| ENSG00000184983 | NDUFA6 | NADH dehydrogenase (ubiquinone) 1 alpha subcomplex, 6, 14kDa | 57 | rs1023500, rs6002655 | chr22:42315744-42689414 | 1.71×10<sup>-9</sup> |
| ENSG00000213593 | TMX2 | thioredoxin-related transmembrane protein 2 | 59 | rs9420 | chr11:57386294-57682294 | 2.24×10<sup>-9</sup> |
| ENSG00000117020 | AKT3 | v-akt murine thymoma viral oncogene homolog 3 | 64 | rs10803138, rs77149735, rs14403, chr1_243881945_I | chr1:243503719-244002945 | 3.73×10<sup>-9</sup> |
| ENSG00000126461 | SCAF1 | SR-related CTD-associated factor 1 | 106 | rs56873913 | chr19:50067499-50135399 | 4.69×10<sup>-8</sup> |
| ENSG00000255408 | PCDHA3 | protocadherin alpha 3 | 108 | chr5_140143664_I | chr5:140023664-140222664 | 4.85×10<sup>-8</sup> |
| ENSG00000204967 | PCDHA4 | protocadherin alpha 4 |  |  |  |  |
| ENSG00000204965 | PCDHA5 | protocadherin alpha 5 |  |  |  |  |
| ENSG00000081842 | PCDHA6 | protocadherin alpha 6 |  |  |  |  |

<sup>a</sup>HUGO Gene Nomenclature Committee ID. <sup>b</sup>Human Genome version 19.

### Polygenic Co-expression Index

To translate *Darkgreen* co-expression into clinical phenotypes, we generated an index predicting *Darkgreen* co-expression based on the genetic background of each individual. We first identified single nucleotide polymorphisms (SNPs) predicting co-expression (co-eQTLs) of the whole module and generated a Polygenic Co-expression Index (PCI (*10*, *14*)). We used a Robust Linear Model to assess allelic dose effects on *Darkgreen ME* (which explained 28% of the variance in the LIBD dataset). The linear model was adjusted for diagnosis, age, sex, RNA integrity (RIN), total RPKM mapped, total RPKM mapped to mitochondrial DNA, and 10 genomic principal components accounting for population stratification. With the aim of increasing our statistical power, we computed a meta-analytic p-value for each SNP based on the effect size in the LIBD and CMC datasets (meta-analytic dataset; overall, 688 subjects). We ranked SNPs based on their meta-analytic p-value and computed several PCIs by adding one SNP at a time (SNPs weights are available in Table S8). Our purpose was to identify an ensemble of SNPs affording prediction of co-expression (correlation between *Darkgreen ME* and PCIs), rather than identifying single genetic variants associated with co-expression per se (although it is noteworthy that the first ranked SNP, rs9836592, would survive Bonferroni correction for multiple comparisons). To determine how many variants should be included in the PCI, we replicated the association between *Darkgreen ME* and PCIs in two additional transcriptomic and genomic datasets (BRAINEAC samples with RIN > 5.5, N = 38; LIBD developmental series samples with RIN ≥ 7.0, N = 93) (*31*, *32*)). The test sets did not affect the model at any stage, because both the *ME* and the weights of the SNPs in the PCI were derived from the training sets. We found that all PCIs including between 6 and 32 SNPs afforded significant predictive capacity in both datasets with an effect size comparable between discovery and replication sets (BRAINEAC: p-value < .05, Fig. 2A-B; LIBD developmental series: p-value < .05; Fig 2A and Fig. S8). Table 3 includes annotations of the first 32 SNPs.

To study translational phenotypes in a clinical population, we performed a meta-analysis of the BRAINEAC and the LIBD developmental series - both test datasets independent of the training sets - to select the most reliable predictors of co-expression. Prediction strength reached a plateau between 14 and 17 SNPs, with no further improvement when more SNPs were added (Fig. 2C). Based on these results, we used the PCIs including 14 to 17 SNPs as predictors of symptom improvement (positive, negative, and general PANSS) in the CATIE clinical trial of antipsychotic efficacy.

**Fig. 2.**
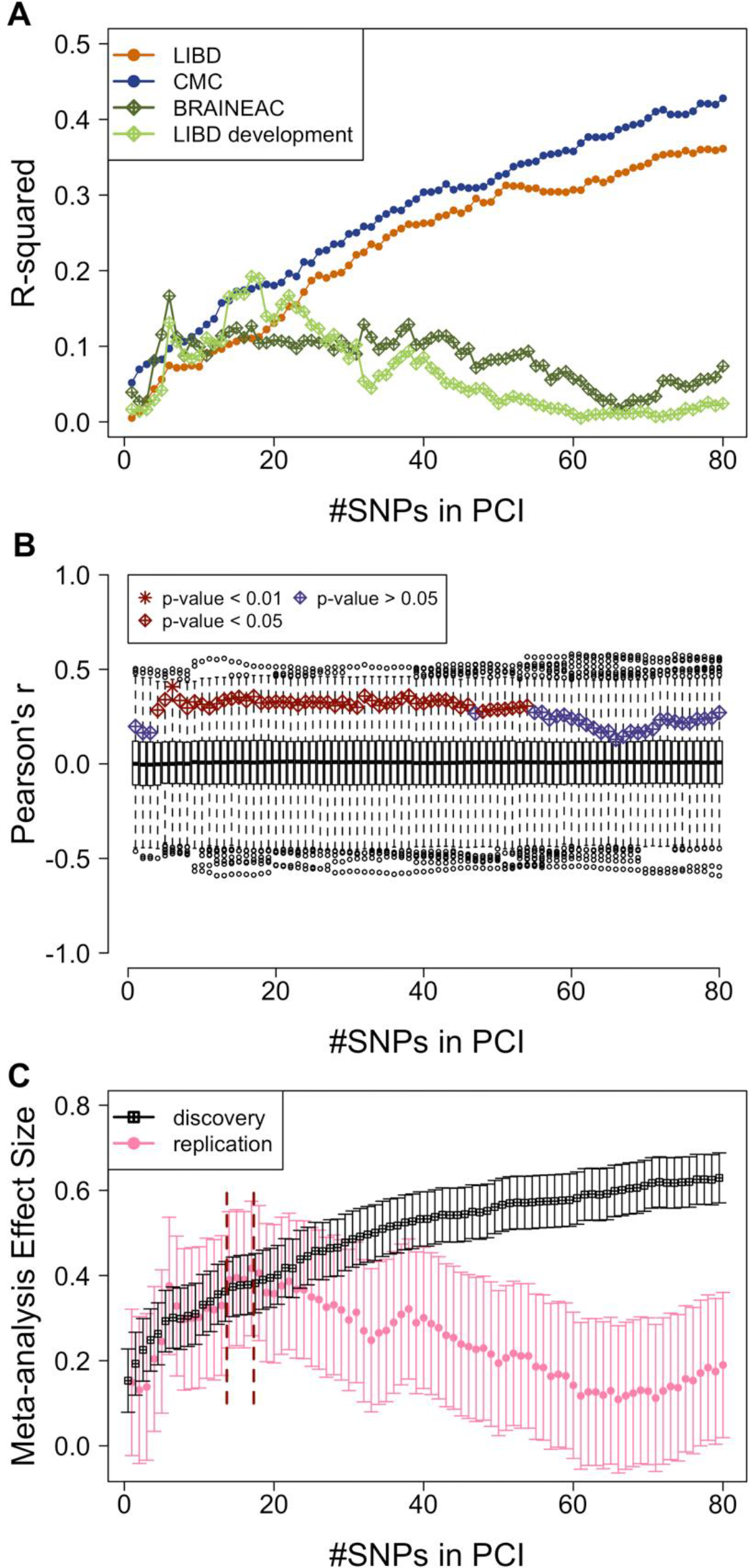
Polygenic Co-expression Index. (**A**) The plot illustrates the variation of the effect size of the correlation between the PCI and the *Darkgreen* module eigengene (Y-axis) for a series of PCIs with incrementally added SNPs. The discovery (LIBD, CMC) and replication datasets (BRAINEAC, LIBD development) are represented with different colors. For increasing number of SNPs included in the PCI (x-axis), the effect size in the discovery sets increases monotonically because of overfitting, while it remains stable and then drops in the replication set, suggesting an optimal signal-to-noise ratio in the replication set between 6 and about 40 SNPs. (**B**) PCI replication. Empirical significance of the correlations between PCIs and Module Eigengene (ME) in the replication set (BRAINEAC). Stars and diamonds display on the y-axis the significance of each *ME*-PCI correlation over an increasing number of SNPs (x-axis). Box plots show the corresponding null distribution of the correlation coefficients when genotypes are permuted (2,000 permutations). Color and shape key in the panel highlight different empirical significance cut-offs. (**C**) Meta-analysis of the effect sizes in the discovery and replication datasets. Dark red vertical dashed lines delimit a plateau in the replication effect sizes between 14 and 17 SNPs. Note that the effect size never increases above the level observed at the 17^th^ SNP.

**Table 3.**
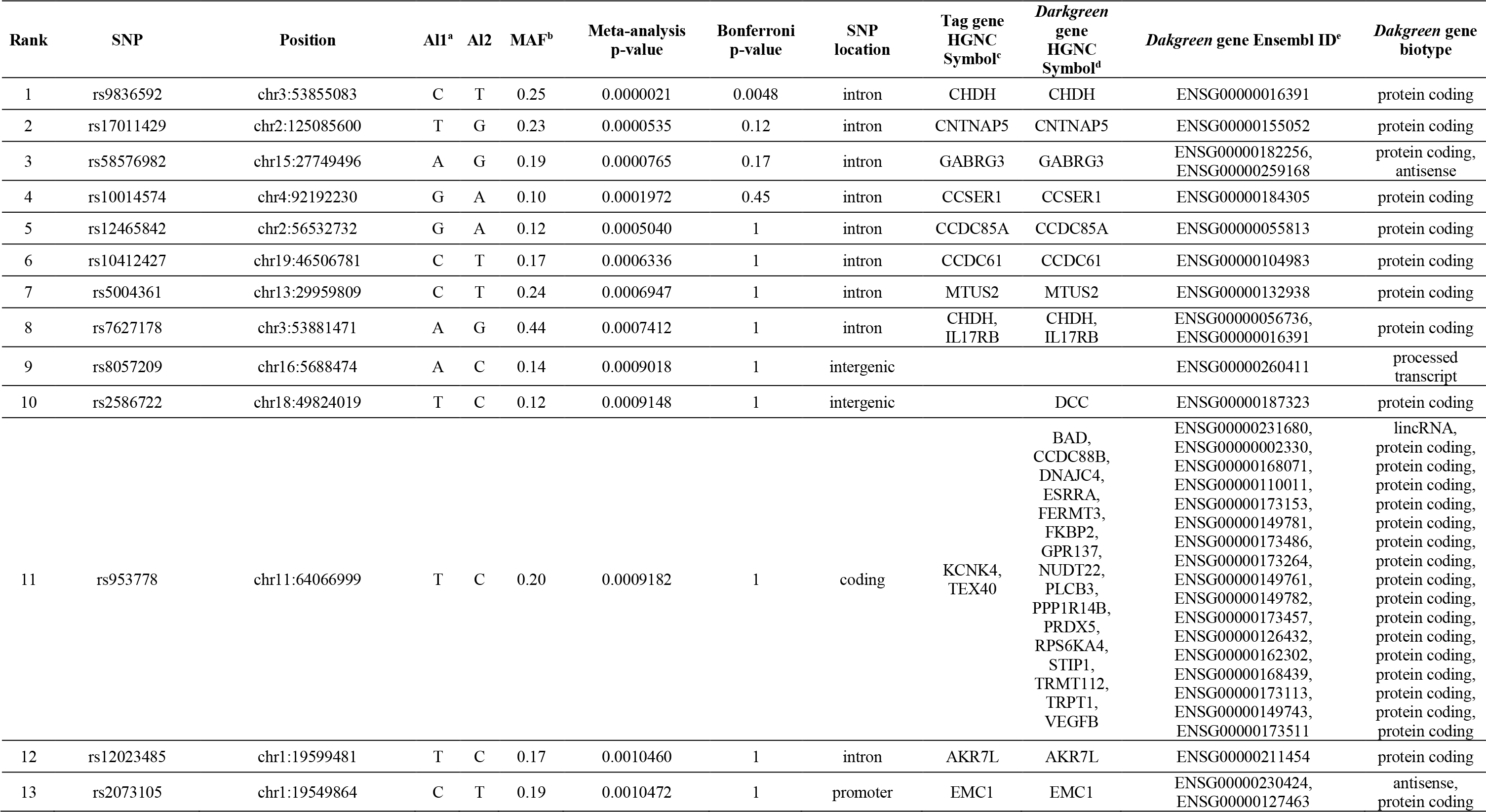
SNP annotations.

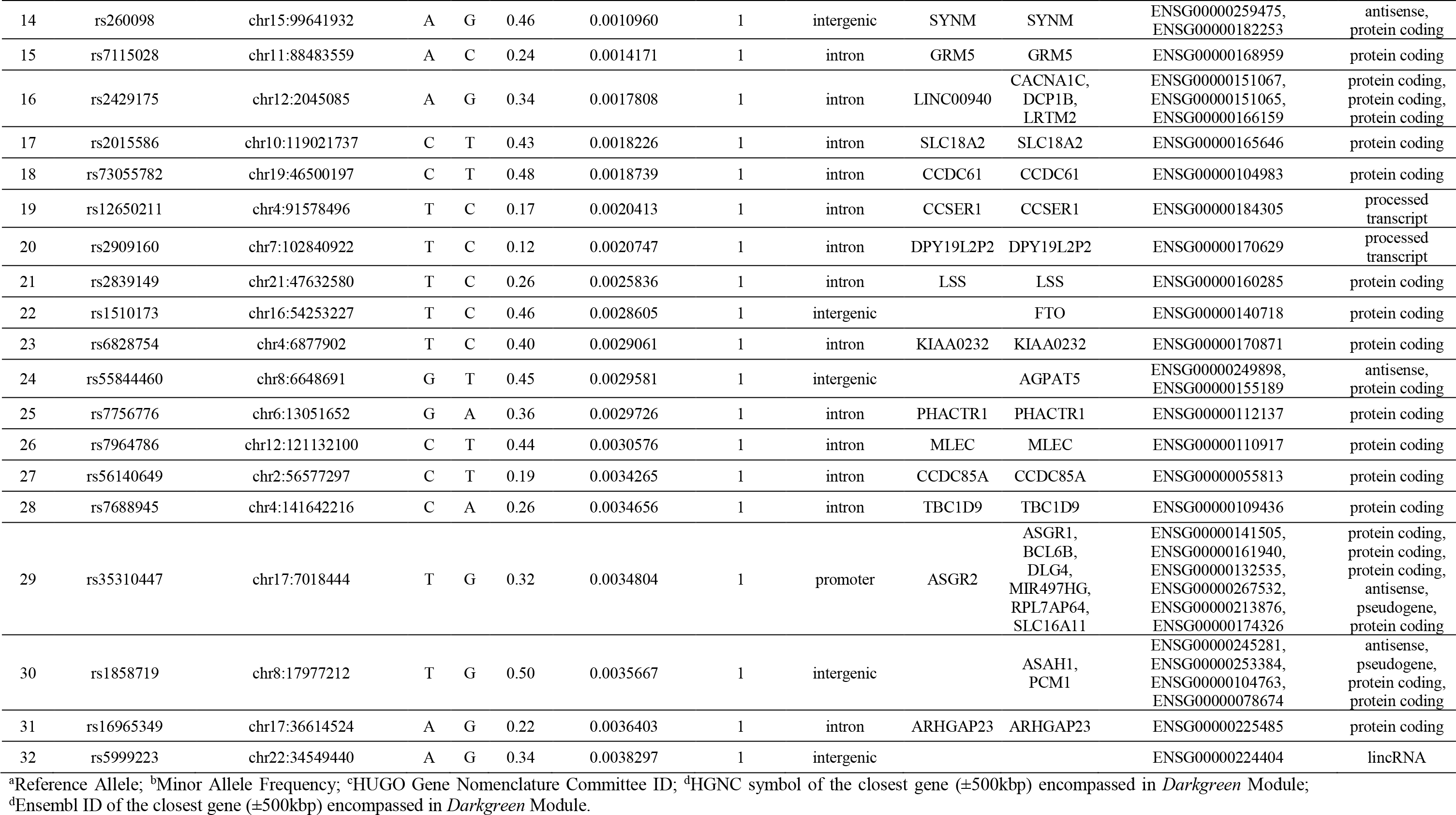

### Clinical study

We focused on patients treated with olanzapine because it showed the best response in the study (*33*) and because we had a replication sample available undergoing the same treatment. The outcome variable was percent change of symptom severity from baseline to one-month follow-up both in CATIE and in UNIBA datasets. We computed a Robust Multiple Regression to assess the association with the PCIs, controlling for age, gender, education level and ancestry (indexed using the first ten genomic principal components). We corrected statistics for multiple comparisons using pACT (*34*). Table 4 illustrates the results. This correction procedure accounts for the high correlation between the predictors and between the dependent variables. We found the most significant relationship between the PCI-16 and positive PANSS improvement (corrected p-value = .033, partial-η^2^ = .061; Fig. 3A), which replicated in the UNIBA independent clinical sample (one-tailed p = .0475, partial-η^2^ = .067; Fig. 3B; Table 4). We assessed the biological significance of this set of 16 SNPs by interrogating Haploreg v. 4.1. Haploreg tests the presence of genetic regulatory elements in a given SNP list (*35, 36*). Our SNP list was specifically enriched for H3K27ac-H3K9ac marks in the dorsolateral prefrontal cortex including Brodmann Areas (BA) 46 and 9 (Bonferroni-corrected p-value = .029). It is worth mentioning that the LIBD RNA sequencing was obtained on BA 46 cortical tissue.

**Fig. 3.**
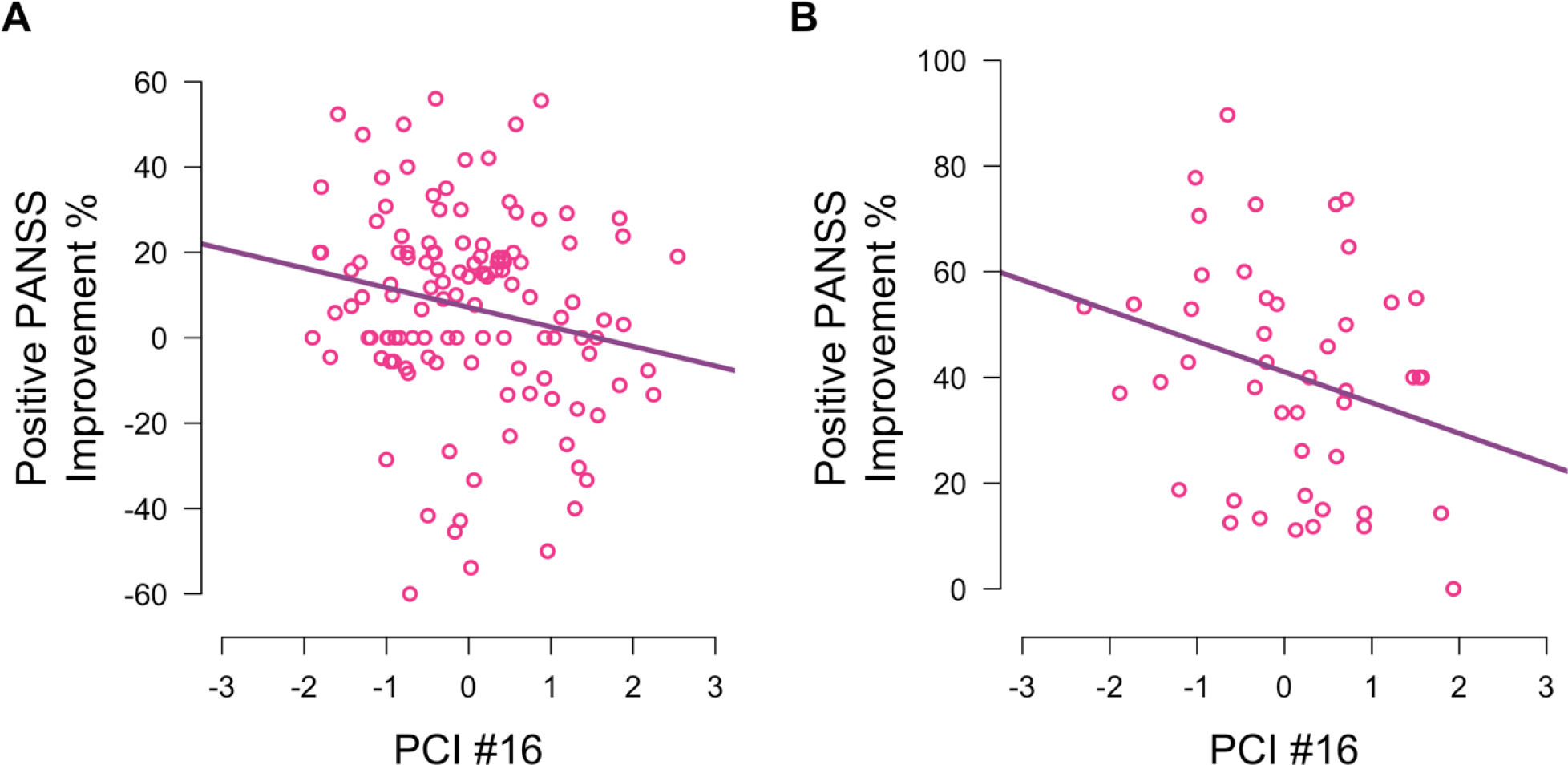
Association between the PCI and clinical outcome. Negative correlation between the PCI with 16 SNPs and symptom improvement in the positive domain of the PANSS (difference between endpoint and baseline relative to baseline, shown on the Y-axis) in the **(A)** CATIE and **(B)** UNIBA cohorts.

**Table 4.** Association between PCIs and positive PANSS early treatment response.

| PANSS subscales | # SNPs in the PCI | CATIE |  |  | UNIBA |  |
| --- | --- | --- | --- | --- | --- | --- |
|  |  | t-value | p-value | Corrected p-value | t-value | One-sided p-value |
| Positive | PCI #14 | -2.11 | .03681 | .134 | -1.93 | .0306 |
|  | PCI #15 | -2.74 | .00714 | .033 | -1.79 | .0403 |
|  | PCI #16 | -2.74 | .00708 | .033 | -1.71 | .0475 |
|  | PCI #17 | -2.58 | .01119 | .049 | -1.60 | .0553 |
| Negative | PCI #14 | 0.59 | .5575 | 1 | 1.14 | .1311 |
|  | PCI #15 | 0.29 | .7750 | 1 | 1.18 | .1218 |
|  | PCI #16 | 0.28 | .7816 | 1 | 1.35 | .0925 |
|  | PCI #17 | 0.22 | .8198 | 1 | 1.42 | .0817 |
| General | PCI #14 | -1.52 | .1301 | .277 | -0.28 | .3889 |
|  | PCI #15 | -1.81 | .0721 | .191 | -0.19 | .4234 |
|  | PCI #16 | -1.74 | .0837 | .208 | -0.24 | .4075 |
|  | PCI #17 | -1.78 | .0771 | .198 | -0.23 | .4106 |

## Discussion

We investigated the convergence of SCZ PGC loci into co-expression networks with the aim of identifying a biological pathway of SCZ risk and regulatory elements associated with gene coexpression that could be translated to the clinic. We identified a gene co-expression module enriched for genes located in risk loci for SCZ. This finding was reproducible, as demonstrated by network preservation and replicated topological overlap in four independent brain gene expression datasets. Module genes were associated with potential gene expression regulation elements. Co-eQTLs identified in 688 subjects were associated with short-term treatment response to olanzapine - a first line antipsychotic - in patients with SCZ. These findings suggest a significant degree of coherence of SCZ risk genes and co-expression partners that might be translated to the clinic.

### Gene co-expression in schizophrenia

In the context of noncoding variation, which characterizes most GWAS significant SNPs and common disorders, gene expression is likely the phenotype closest to DNA in which inter-individual differences can be directly associated with genetic variation. The multifold preservation of the network is important because one may expect that gene co-expression in patients with SCZ may be confounded by state-related factors such as pharmacological treatment; instead, our results demonstrate that such state-related factors did not dominate the topology of the network, which was replicated in three independent datasets of non-psychiatric individuals of various ages totaling 227 subjects. Therefore, it is unlikely that our results are biased because of the use of data from patients. Moreover, we failed to associate the gene-gene relationships within *Darkgreen* with smoking or antipsychotic medication (90 patients were treated and 50 showed no evidence of treatment with antipsychotics), though these phenomenological factors are poorly quantified in *post-mortem* tissue. Notably, it is difficult to conclusively rule out the effect of antipsychotic medication because the possible confounding effects of medication may depend on the specific antipsychotic administered and on the dosage.

Jaffe and coworkers (*12*) have suggested that preprocessing RNA data controlling for hidden RNA quality is a key factor affecting the inferences drawn from transcriptome studies and the topology of the network we report here holds also when preprocessing data with the conservative approach they described (Fig. S11). In summary, the network we identified and validated in the largest sample tested to date (including data from overall 915 *post-mortem* samples) is robust in terms of reproducibility and highlights gene-gene relationships revealing non-random clustering of SCZ risk genes.

### The schizophrenia risk co-expression module

Gene ontology analysis revealed involvement of *Darkgreen* genes in cell-cell adhesion, a biological process previously associated with risk for SCZ and bipolar disorder (*37*, *38*). It should be noted that we selected genes expressed in the brain, whereas ontologies were not filtered in the same way. This implies that our approach was conservative and more biological functions than currently detected may be shared by these genes. Interestingly, the same gene ontology characterized differentially expressed genes in induced pluripotent stem cell-derived differentiated neurons, in a recent study comparing populations of monozygotic twins with discordant response to clozapine in treatment-resistant SCZ (*39*). Taken together, both findings highlight the potential importance of the genes co-expressed in *Darkgreen* for the physiology of olanzapine and clozapine, two atypical antipsychotics. *Darkgreen* included also genes coding for proteins involved in synaptic transmission mediated by serotonin, glutamate and GABA (*HTR1F*, *GRM5*, *GABRB1*, *GABRG3*), or involved in neural excitability (*KCNH1*, *KCNA3*, *KCNH7*, *KCNH5*), along with *CACNA1C*, a risk gene for SCZ and bipolar disorder supported by multiple lines of evidence (*40*–*44*). The functions of the genes in *Darkgreen* are consistent with previous pathway analyses of SCZ risk (*45*) and enhance the biological plausibility that the coregulation of this module has functional relevance.

Although co-expression does not necessarily imply gene co-regulation, it is noteworthy that the 13 PGC hits of *Darkgreen* are distributed across 10 different loci, rather than encompassing a single locus that is co-transcripted because of genetic proximity (*48*) and the significant preservation of *Darkgreen* topology in very young subjects is consistent with the hypothesis that SCZ risk genes are coordinated by processes relevant to neurodevelopmental trajectories.

### Genetic variants associated with co-expression of schizophrenia risk genes

Since it is not possible to directly assess gene expression in the living human brain, it is of interest to translate models of gene co-expression into genetic variants (co-eQTLs) which index co-expression in living individuals. The co-eQTLs detected here merit further investigation as potential indicators of loci affected by genetic regulatory elements associated with positive symptoms and their clinical course. For example, the first ranked SNP, rs9836592, has been associated with risk for bipolar disorder (*49*), another disorder frequently treated with antipsychotic drugs such as olanzapine. Furthermore, this SNP has been already associated with the regulation of gene expression (*49*) and is an eQTL for *CACNA7D* (*9*). Moreover, the entire set of 16 SNPs was enriched for histone acetylation marks. Previous evidence supports the relevance of histone modification pathways to SCZ (*45*) and the specific role of H3K27ac markers in autism (*50*) a neurodevelopmental disorder sharing some genetic risk with SCZ (*57*). The clinical evidence obtained in two independent samples supports the functional role of these SNPs in the clinical treatment of SCZ.

### Clinical translation of transcriptome data mining

We found that the PCI computed using the genetic variants above described was reproducibly associated with treatment response to olanzapine. On the one hand, this finding suggests that the 13 PGC hit genes co-expressed in *Darkgreen* are candidates within their loci for mechanistic interpretations of response to treatment. On the other hand, the PCI indexes a wider group of genes, going beyond the 13 PGC hits, suggesting a broader transcriptomic landscape of risk and more relevant here, of the biology of treatment response. Such landscape stratifies patients with SCZ in terms of treatment response even though *Darkgreen* co-expression is not reproducibly associated with diagnosis and the PCI variants per se are not associated with diagnosis. Another implication of the present findings is that antipsychotic efficacy may involve many more genes than those coding for the traditional targets, e.g., dopamine and serotonin antagonism, and may depend on the convergence in terms of genetic regulation of multiple neural transmission systems, including glutamate and GABA receptors, as well as calcium and potassium channels. This possibility is implicit in the fact that dopamine and serotonin are engaged in tuning glutamate and GABA neuronal activity in cortex (*52*, *53*).

Our findings further suggest a link of SCZ risk loci and their molecular interactors with inter-individual variation in response to treatment with olanzapine selectively in terms of positive symptoms domain, despite the differences between the clinical datasets we used. The current evidence is limited by the relatively restricted sample size in the clinical groups (total N=167) and by the modest size of the clinical effects. Therefore, this evidence awaits further independent replications in larger clinical samples. However, this clinical translation is promising with respect to the feasibility of patient stratification based on biological measures, in line with dimensional views of the diagnosis of SCZ (*57*).

This study demonstrates the potential for co-expression genetic studies to be translated in the clinic. However, several limitations suggest caution. First, while WGCNA is a flexible and extensively used tool, gene co-expression network analyses can be implemented with different methodological nuancing. For example, reproducible gene-gene relationships can be reflected in different gene clustering across datasets and studies. Second, a large portion of the variance in treatment response remains unexplained (> 90%), suggesting the potential role of other factors not assessed here. Large datasets including longitudinal clinical information, genome-wide genotyping, along with brain imaging data and environmental variables, may bring us closer to the clinical utility of this work (*58*). Third, the role of potential regulators of gene co-expression requires biological evidence to offer mechanistic explanation of how their targets are related with response to olanzapine. Addressing these limitations will be necessary steps to more routinely apply genetic screening in the clinic.

Nevertheless, this work demonstrates that a proportion of SCZ risk genes converge into gene coexpression networks and provides information on potentially relevant molecules implicated in this process. The findings offer a proof of concept that translation of genetic risk into clinical information requires the study of multiple levels of biological organization, starting from the very beginning of the information flow from DNA to phenotypes, i.e., gene expression.

## Material and Methods

### Study design

Table 1 summarizes the demographic data and relative statistics for the subjects included in all experiments. For the co-expression network study, we used RNA sequencing data from the LIBD (*12*) and from the CMC (*8*) *post-mortem* series for a transcriptome-wide WGCNA (*13*). Both datasets included *post-mortem* mRNA expression levels of HCs and patients with SCZ in the human prefrontal cortex, whereas the three additional datasets used for replication included only non-psychiatric individuals (*31*, *59*). Additionally, the LIBD dataset included toxicological tests performed on frozen *post-mortem* tissue long after death. Smoking habit was assessed based on nicotine and cotinine quantification, as well as on reports from familiars. Drug consumption assessment, particularly regarding antipsychotics and antidepressants, has been recorded as a yes/no variable. Permission to use *post-mortem* brain materials was obtained by the next of kin (see the original reports for further information). We selected subsets of individuals in the LIBD and CMC datasets to match possible confounding variables across the datasets as closely as possible. Therefore, we included samples with age ≥ 17 years of Caucasian or African American ancestry, RNA integrity number (RIN) ≥ 7.0. We used χ^2^ tests to assess the effects of gender, ethnicity and diagnosis between datasets and a two-sample t-test to assess the effect of age.

In the clinical studies, all participants provided written informed consent following the guidelines of the Declaration of Helsinki after receiving a complete description of the study. Protocols and procedures were approved by the ethics committee of the University of Bari (UNIBA) and by the institutional review board of each clinical site involved in the CATIE program. Diagnosis of Schizophrenia was established via Structured Clinical Interview for DSM-IV-TR (SCID). Symptom severity was assessed with Positive and Negative Syndrome Scale (PANSS) (*60*) at study entry and at several follow-up visits. The first clinical cohort included patients recruited in CATIE study by the NIMH and treated with olanzapine (N = 121) (*33*). The second cohort included 46 patients recruited from the region of Apulia, Italy, also treated with olanzapine in monotherapy (*16*). Study protocols and exclusion criteria are available in SM Materials and Methods.

### Co-expression Network of human prefrontal cortex

We processed RNA sequencing raw data as previously described (*63*). RUV capitalizes on the putatively low physiological variation of housekeeping genes (HK). Therefore, variation in HK expression may reflect more closely systematic noise than inter-individual variability. This version of RUV was specifically designed to correct the signal prior to WGCNA (SM Material and Methods).

WGCNA (*13*, *64*) uses gene-gene Pearson’s correlation indices as a continuous, i.e., weighted, measure of gene-gene relationships. We computed unsigned networks, i.e., negatively correlated genes are considered connected rather than non-connected (*65*). The correlation matrix was transformed into an adjacency matrix by raising Pearson’s coefficients to a positive exponent, β, which is chosen to meet the “scale-free” power law connectivity distribution. Scale invariance is widely considered a common organization feature of cellular functions (*66*). A hierarchical clustering method was then used to group genes into clusters, called “modules” (SM Materials and Methods). Colors were used to arbitrarily label co-expression modules, with the “grey” module representing genes that did not cluster into any particular module. Co-expression was summarized by the *ME*, the first principal component of the expression of genes in any given module. A unique *ME* was computed for each module.

First, we computed separate co-expression networks for patients with SCZ and HCs within each of the two datasets. To identify possible differences in network topology between patients and controls, we employed the same β value for all datasets, because this parameter affects mean network connectivity (β = 6 was the minimum value that satisfied the scale invariance criterion for all datasets, which fits well with the authors’ suggestions for unsigned networks; for signed networks higher exponents are generally needed, e.g., β = 12). We used the methods described by Langfelder et al.(*18*) and by Johnson et al.(*26*) to compare graph properties using permutation approaches. These procedures are complementary because the first relies on evaluation of graph parameters, while the second entails an empirical, parameter-free test. The preservation technique published by Langfelder and coworkers (*18*) uses connectivity and density to derive a summary score that characterizes optimal preservation with Z ≥ 10, partial preservation with 2 ≤ Z < 10, and no preservation with Z < 2 (1,000 permutations). The technique developed by Johnson et al. (*26*), instead, assesses whether the topological relationships between genes in the second dataset mirror those of the first dataset at a level greater than chance. Therefore, for each module we computed the median of its topological overlap matrix and compared this value against the null distribution of medians computed on random modules of identical size. We used 10,000 re-samplings and a threshold for replication significance of empirical p-value < .001.

Network statistics showed strong preservation between HCs and patients with SCZ within each dataset. Moreover, we used Wilcoxon signed rank test to demonstrate that preservation statistics (Z-values) were greater between groups within the same dataset than between the same group across the two datasets (SM Results, Fig. S1). Based on these results, we adopted an alternative approach. We pooled data from patients with SCZ and HCs and identified one network for the LIBD and one for the CMC datasets, allowing greater statistical power for the next steps of the analysis. All the following analyses used the LIBD network with pooled patients with SCZ and HCs as the reference set. The minimum value of β that satisfied scale invariance criterion both in the LIBD and in the CMC datasets was 5. This network was comprised of 43 modules, with 6,706 transcripts falling in the grey module, i.e., not clustered (Data file S1). These modules were strongly preserved in CMC (Fig. 1a-b, Table S1). Importantly, the “gold” module, i.e., a random module whose size was defined as equal to the median of the sizes of all modules, showed the lowest Z preservation statistic (Fig 1a). We cross-checked preservation using CMC as the reference (57 modules, grey: 7,228 transcripts; also in this case, all modules had Z ≥ 2; Table S2).

We assessed the association of the LIBD Module Eigengenes (*MEs*_*LIBD*_) with case-control status (i.e., HCs vs. patients with SCZ) with a Robust Linear Model with the *lmRob* function of the *robust* R package. We used Bonferroni correction for multiple comparisons (corrected p-value < .05). We introduced observed demographics, RNA quality feature and RNA sequencing coverage as covariates since they may potentially affect gene expression measures. The model accounted for age, gender, RIN, total reads mapped, total reads mapped at mitochondrial DNA and 10 genomic ancestries as covariates to account for potential genetic stratification (SM Materials and Methods). Then, we replicated the findings in the CMC dataset. In order to obtain factor scores in the CMC network (*MEs*_*CMC*_) for each corresponding LIBD module, we computed the factor loadings for each *ME*_*LIBD*_. Factor loadings express the weighted contribution of each gene in the module to the *ME*. Then, we projected factor loadings into the corresponding CMC gene expression data to obtain projected-MEsCMC. Thus, we evaluated the replication of the association between co-expression (*projected-ME*_*CMC*_) and case-control status in the CMC dataset (p-value < .05).

In order to address the effect of potential confounders on the identified network, we employed Robust Linear Models to assess the association between each *ME*_*LIBD*_ and nicotine, cotinine and smoking status separately. Moreover, we evaluated the association between each *ME*_*LIBD*_ with antipsychotics and antidepressants in the SCZ group (α = .1; Data file S2).

### Co-expression Network replication

We used several datasets of *post-mortem* brain samples to validate the LIBD modules through the above-mentioned permutation procedure (*26*).

i. The LIBD Developmental Series (spanning ages from fetal to adolescent);
ii. The CMC dataset (as already described);
iii. The BRAINEAC Frontal Cortex dataset (*31*);
iv. The GTEx Brain Cortex dataset (*59, 67*);
v. The GTEx Frontal Cortex Brodmann Area 9 dataset (*59, 67*).

Datasets i-ii) were pre-processed as described in the pre-processing section. Dataset iii) is publicly available at http://www.braineac.org/. Microarray expression data were downloaded and pre-processed through RUV tools, selecting the same parameters used for the LIBD and CMC datasets (k = 5). Datasets iv-v) are available at https://www.gtexportal.org/home/. RNA sequencing data have been downloaded in the already pre-processed release format (GTEx Analysis V6p), with the aim to test for module replication regardless of the pre-processing pipeline.

### Prioritization of modules relevant for SCZ

The enrichment analysis is used to characterize the functional profile of gene sets identified *a priori*. It consists in identifying over-represented gene classes within another gene set.

1. We investigated the overlap between the LIBD modules and genetic association with SCZ (*1*). We referred to genes identified by the PGC study (n = 310 genes were included in the network based on transcript expression levels, Table S4) and used a hypergeometric test to assess the significance of the over-representation in each module. We selected the modules surviving Bonferroni correction for multiple comparisons (number of modules = 43, p-value = .05/43 = .00116). Moreover, we conducted hypergeometric tests at multiple levels of PGC loci expansion (from ±50 kbp to ±10Mbp) to investigate the range of the gene-gene interactions potentially involved in the convergence of SCZ risk genes. We used the biomaRt R package (*68*) to select protein coding genes located within the expanded PGC loci. Finally, we derived an empirical p-value through a permutation approach (p < .001, SM Materials and Methods). Since restricting the analysis to protein coding genes may bias results because network analysis encompasses different gene biotypes (protein-coding and non-protein-coding), we repeated the same analysis also including all the genes located in the expanded PGC loci, regardless of gene biotype (Fig. S3).
2. In addition, we explored the enrichment for common SCZ variants. We used summary statistics of 9.4 million SNPs from the largest GWAS in SCZ by PGC (*1*) publically available (http://www.med.unc.edu/pgc/results-and-downloads) and excluded the MHC region on chromosome 6 because high LD in this locus could bias gene set enrichment statistics as already suggested by other authors (*45*). We used Multi-marker Analysis of GenoMic Annotation (MAGMA)(*22*) to perform a gene-set competitive enrichment analysis adjusted for confounding variables (SI Materials and Methods). The significance threshold was set at the nominal p-value < .05 because we were only interested in modules that already showed a significant overrepresentation of SCZ risk genes.

#### Functional enrichment analyses

We used Amigo2 (http://amigo2.geneontology.org/amigo, Gene Ontology database released 2017-0609) online available tools to perform functional enrichment analyses of the *Darkgreen* module, which was selected based on the overrepresentation of SCZ risk genes. We listed *Darkgreen* protein coding genes and performed online automatic searches in the Gene Ontology Database Released on 2017-0629 with the PANTHER Overrepresentation Test (release 2017-04-13). Furthermore, we used Specific Expression Analysis (SEA) software [http://genetics.wustl.edu/jdlab/csea-tool-2/] (*23*) to track cell- and tissue-specific expression pattern during neurodevelopment (Fig. S4).

#### Enrichment analysis of Transcription Factor Binding Sites

We used Pscan, a freeware web interface (http://159.149.160.88/pscan/)(*27*) to scan promoter regions of our co-expressed genes looking for binding specificity of known Transcription Factors (TF). We referred to the JASPAR 2016 (*69*) database of TF binding profiles and defined the promoter regions spanning 1,000 bp upstream the transcription starting site by selecting these options from the web interface. We scanned 472 different TF binding domains. We considered statistically significant TFs surviving Bonferroni correction for multiple comparisons (corrected p-value < .05). Then, we explored the contribution of single genes to the selected TFs and reported SCZ risk genes contained in *Darkgreen* related with the TF more strongly than genome-wide average for the same TF (*27*)). Finally, to explore the specificity of our findings, we evaluated the enrichment of all the other modules and reported corrected p-values (Fig. S5).

#### Micro-RNA target prediction

We investigated the overlap between SCZ related miRNA targetomes (*29*) and *Darkgreen*, with the purpose to identify specific regulatory elements of co-expressed genes. We used four miRNA target repositories to obtain different lists of targets for each miRNA family (i. TargetScan v7.1, http://www.targetscan.org/vert71/ (*70*); ii) MirTarget, http://www.mirdb.org/(*71*); TargetMiner, http://www.isical.ac.in/~bioinfo_miu/targetminer20.htm(*72*) and TarBase V7.0(*73*)). Then, we performed a hypergeometric test for over-representation of miRNA targets in *Darkgreen* and combined p-values with sum-log Fisher’s method across different lists for each miRNA family. The corrected significance threshold for the combined p-values was set to p-value = .00125, after having applied Bonferroni correction (10 miRNA families tested times 4 tools used). We inspected targetomes overlapping with *Darkgreen* and reported SCZ risk genes available in at least one gene list (Table S7). Finally, we investigated the specificity of these enrichments by extending the same analysis to all the LIBD modules and miRNA families (Data file S3). We expected that each miRNA would be associated with only few modules. We showed results corrected for multiple comparisons (Bonferroni rule, number of modules = 43; Fig. S6).

#### Overlap with genes regulated by haloperidol

We investigated the overlap between putative antipsychotics target genes and *Darkgreen* as well as all other modules. We used lists of genes differentially expressed (DEG) between haloperidol-treated mice and the control mice (*30*). We used lists of DEG at q-value < .05 in the striatum and in the whole brain of mice (*30*). Moreover, we separately tested up- and down-regulated genes. We employed biomaRt R package (*68*) to convert mouse genes into human orthologs. We performed hypergeometric test for over-representation of haloperidol targets in network modules and used Bonferroni correction for multiple comparisons (number of modules = 43; Fig. S7).

#### Meta-analysis of co-expression quantitative trait loci

SNP genotyping procedures and genotype imputation have been described previously for LIBD (*9*), CMC (*8*), BRAINEAC (*31*), CATIE (*33*) and UNIBA (*10*) subjects (also see SM Materials and Methods). We selected SNPs in the genes encompassed in *Darkgreen*, expanded by 100 kbp up- and down-stream gene start and end, consistent with previous studies (*10*, *14*). We employed a relatively conservative extension of the genes because with larger flanks, e.g., 500 kbp to 1 mbp, the SNP sets would largely overlap between modules. We selected SNPs with MAF ≥ 0.1 because the sample size was too limited to investigate uncommon variants and pooled minor allele carriers when MAF ≤ 0.15 to avoid biasing estimations of population variance with small genotypic groups. These filters resulted in 52,198 SNPs available in both the LIBD and the CMC datasets that we selected for further analyses.

We aimed to identify an ensemble of SNPs that, together, could predict gene co-expression (co-eQTLs). The biological plausibility of co-eQTLs is supported by findings that only 30% of mRNA expression heritability is associated with cis-active elements (*74*), suggesting a role of distant regulatory elements and possibly *trans*-elements in heritable mRNA expression. With this purpose, we investigated the association between the *Darkgreen* co-expression module summarized by the *ME*-*Darkgreen* (see section 2.3) and SNP allelic dosage. We used a Robust Linear Model to estimate the effect of the SNP allelic dosage separately in the LIBD and the CMC datasets with the *lmRob* function of the *robust* R package. We included diagnosis (HCs vs. patients with SCZ), age, gender, RIN, total count of mapped reads, total count of mitochondrial mapped reads and 10 ancestries as covariates. Notably, co-varying for diagnosis allowed us to detect markers of co-expression valid both in patients and controls rather than risk markers for SCZ. Finally, we performed a fixed-effect meta-analysis over the two datasets with the *rma.uni* function of the *metaphor* R package, using partial correlation coefficients of allelic dosage as an estimate of effect size. Then, we ranked SNPs according to their meta-analytic p-value.

Following previous work on polygenic summaries of additive genetic effects (*1*, *10*), we restricted the analysis to independent SNPs. In this perspective, we evaluated pair-wise R^2^ between SNPs within 250 kbp. We considered two SNPs independent when R^2^ < 0.1 (*1*). We then performed a priority LD pruning by iteratively discarding the SNP with the weaker association. We used this procedure to enrich our selection for relevant variants (for further applications of a similar procedure see http://prioritypruner.sourceforge.net/documentation.html). The final selection included 2,266 tagging SNPs with negligible residual interdependence. We used the top 100 ranked co-eQTLs for the PCI computation.

#### Polygenic Co-expression Index

We employed a previously published procedure based on Signal Detection Theory to assign weights (A’) to each SNP genotype (*10*, *14*) (SM Materials and Methods). For each genotypic population of each of the 100 top-ranked SNPs, we computed the A’ weights separately in the LIBD and the CMC datasets. Then, we averaged the weights across the two datasets (Table S8). We defined the PCI as the average of A’ values corresponding to all the genotypes of each subject. In this way, the PCI could be interpreted as the genetically indexed inter-individual variability associated with gene co-expression measured by Darkgreen Module Eigengene (*ME-Darkgreen*). The PCI is positively correlated with *ME-Darkgreen* and is not confounded by ethnicity (SM Materials and Methods and Fig. S9).

A relevant issue is how many SNPs need to be included in the PCI. Including too few SNPs may not afford sufficient predictive power, while too many SNPs may yield overfitting effects on the positive correlation between the PCI and the *ME-Darkgreen*. To identify a SNP ensemble with significant predictive power, we computed 100 different PCIs with an increasing number of SNPs (the first PCI included just the first ranked co-eQTL, the second PCI included the first and the second co-eQTL, and so on up to the 100^th^ co-eQTL) and assessed the Pearson’s correlations between the PCIs and the *ME-Darkgreen* both in the LIBD and the CMC datasets. In case of overfitting, the effect size of the correlation PCIs-*ME* in the discovery sets should monotonically increase when more SNPs are added, whereas the effect size in the replication datasets should reach a plateau and then decrease (Fig. 2A). We assessed the statistical significance of the PCI-*ME* correlation in the two independent replication sets also via a permutation approach (p-value < .05, SM Materials and Methods).

Then, we performed a fixed-effect meta-analysis separately on the discovery and replication datasets. In this way, we estimated global replication effect sizes using PCI-*ME* correlation coefficients (Fig. 2C). In order to identify the best set of predictors for the clinical study, we selected PCIs based on the largest replication effect size. We started to include PCIs at the beginning of the plateau and stopped when the effect size reached the absolute maximum and then started to decline (a possible effect of overfitting; Fig. 2C).

#### Clinical study

We used two samples of patients with SCZ treated with olanzapine to assess the association between the PCIs and the clinical outcome measured with the PANSS. Clinical outcome was defined as the difference between baseline and early clinical response (one month) relative to baseline symptoms in PANSS sub-scales and total scores. Patients were genome-wide genotyped (SM Materials and Methods) and SNP genotypes were used to compute PCIs for each patient. We tested the association between clinical outcome and PCIs through a Robust Linear Model using age, gender, education level and ten genomic PCs as nuisance covariates. The CATIE cohort was used as discovery sample and results were corrected for multiple comparisons, i.e. the multiple clinical subscales and PCIs we tested (corrected p-value < .05). Due to the high correlation among the set of predictors - the PCIs - and among the set of outcomes, we used an appropriate procedure for p-values adjustment of multiple correlated tests (*34*). We selected the best model and replicated the association in the UNIBA cohort (one-tailed p-value < .05). We reported the effect size as partial η^2^.

Finally, to assess the biological significance of the SNPs encompassed in the PCI, we submitted the list and the selected variants in full linkage disequilibrium with them to HaploReg 4.1(*36*) selecting American ancestry and all four epigenome sources. Haploreg is a repository of genetic regulatory elements across multiple tissues according to previous genomic studies (*35*, *36*). Since the reference network was identified in dorsolateral prefrontal cortex, we specifically interrogated this brain region including BA46 and 9. Finally, we computed the statistics for overrepresentation of regulatory elements (Bonferroni-corrected p-value < .05).

## Supplementary Materials

### Material and Methods

#### Results

Fig. S1. Results of intra-dataset preservations (Langfelder method).

Fig. S2. Preservation of CMC network in the LIBD dataset (Langfelder method).

Fig. S3. *Darkgreen* module enrichment for schizophrenia risk genes (all gene biotypes).

Fig. S4. Specific Expression Analysis (SEA).

Fig. S5. Transcription factor binding sites enrichment analysis.

Fig. S6. MicroRNA targets prediction.

Fig. S7. Overlap of LIBD modules with haloperidol targets.

Fig. S8. PCIs replication in LIBD developmental ages set.

Fig. S9. Association between PCIs and *Darkgreen* ME separately in Caucasian and African-American subjects.

Fig. S10. RUV pre-processing.

Fig. S11. Preservation of the LIBD network in LIBD dataset with an alternative preprocess pipeline (Langfelder method).

Table S1. The LIBD network: module replication in the CMC dataset and enrichment statistics for schizophrenia risk.

Table S2. The CMC network: module replication in the LIBD dataset.

Table S3. The LIBD network: association between module eigengenes (MEs) and diagnosis (HC vs. SCZ).

Table S4. Genes in the PGC list included in the network.

Table S5. Chart of *Darkgreen* genes and connectivity statistics.

Table S6. The LIBD network: module replication (empirical p-values).

Table S7. Overrepresentation of miRNA targetomes in *Darkgreen* module.

Table S8. SNP annotations: A’ weights of SNP genotypes used for PCI computation.

Data file S1. (Microsoft Excel format). Chart of LIBD network genes, module labels assignments and connectivity statistics.

Data file S2. (Microsoft Excel format). Chart of p-values: association between LIBD module eigengenes and biological confounders.

Data file S3. (Microsoft Excel format). Chart of combined p-values: enrichment of LIBD modules for miRNA targetomes.

## Acknowledgments

This article was based on results from the Clinical Antipsychotic Trials of Intervention Effectiveness (CATIE) project supported with Federal funds from the National Institute of Mental Health (NIMH) under contract NO1 MH90001. The project was carried out by principal investigators from the University of North Carolina, Duke University, the University of Southern California, the University of Rochester, and Yale University in association with Quintiles, Inc., and the program staff of the Division of Interventions and Services Research of the NIMH and investigators from 84 sites in the United States. AstraZeneca Pharmaceuticals LP, Bristol-Myers Squibb Company, Forest Pharmaceuticals, Inc., Janssen Pharmaceutica Products, L.P., Eli Lilly and Company, Otsuka Pharmaceutical Co., Ltd., Pfizer Inc., and Zenith Goldline Pharmaceuticals, Inc., provided medications for the studies. CMC data were generously provided to GP by the NIMH and CommonMind Consortium. We gratefully acknowledge the work by Prof. Roberto Bellotti, Dr. Alfonso Monaco (Department of Physics - University of Bari Aldo Moro), Marco Zezza, Leonardo Sportelli and Elisabetta Volpe (Department of Basic Medical Science, Neuroscience, and Sense Organs - University of Bari Aldo Moro), who contributed to data analysis. We are also in debt to Dr. Gianluca Ursini, Dr. Richard Straub, and Dr. Venkata S. Mattay (Lieber Institute for Brain Development) for insightful discussions on the procedures employed.

## Author contributions

GP, PDC, DRW, and AB designed the study; TMH, JEK, GB, DRW, AR, and AB were involved in data collection; PDC, AEJ, MP, QC analyzed the data; GP, PDC, AEJ, DRW, and AB interpreted the data; GP, PDC, and AB wrote the first draft of the manuscript; all authors revised and approved the manuscript.

## References and notes

1. C. Schizophrenia Working Group of the Psychiatric Genomics, Biological insights from 108 schizophrenia-associated genetic loci. Nature 511, 421–427 (2014).

2. K. S. Kendler, What psychiatric genetics has taught us about the nature of psychiatric illness and what is left to learn. Molecular psychiatry 18, 1058–1066 (2013).

3. N. C. Hettige, C. B. Cole, S. Khalid, V. De Luca, Polygenic risk score prediction of antipsychotic dosage in schizophrenia. Schizophrenia research 170, 265–270 (2016).

4. T. Wimberley, C. Gasse, S. M. Meier, E. Agerbo, J. H. MacCabe, H. T. Horsdal, Polygenic Risk Score for Schizophrenia and Treatment-Resistant Schizophrenia. Schizophrenia bulletin, (2017).

5. E. A. Boyle, Y. I. Li, J. K. Pritchard, An Expanded View of Complex Traits: From Polygenic to Omnigenic. Cell 169, 1177–1186 (2017).

6. P. J. Harrison, D. R. Weinberger, Schizophrenia genes, gene expression, and neuropathology: on the matter of their convergence. Molecular psychiatry 10, 40–68; image 45 (2005).

7. C. Gaiteri, Y. Ding, B. French, G. C. Tseng, E. Sibille, Beyond modules and hubs: the potential of gene coexpression networks for investigating molecular mechanisms of complex brain disorders. Genes, brain, and behavior 13, 13–24 (2014).

8. M. Fromer, P. Roussos, S. K. Sieberts, J. S. Johnson, D. H. Kavanagh, T. M. Perumal, D. M. Ruderfer, E. C. Oh, A. Topol, H. R. Shah, L. L. Klei, R. Kramer, D. Pinto, Z. H. Gumus, A. E. Cicek, K. K. Dang, A. Browne, C. Lu, L. Xie, B. Readhead, E. A. Stahl, J. Xiao, M. Parvizi, T. Hamamsy, J. F. Fullard, Y. C. Wang, M. C. Mahajan, J. M. Derry, J. T. Dudley, S. E. Hemby, B. A. Logsdon, K. Talbot, T. Raj, D. A. Bennett, P. L. De Jager, J. Zhu, B. Zhang, P. F. Sullivan, A. Chess, S. M. Purcell, L. A. Shinobu, L. M. Mangravite, H. Toyoshiba, R. E. Gur, C. G. Hahn, D. A. Lewis, V. Haroutunian, M. A. Peters, B. K. Lipska, J. D. Buxbaum, E. E. Schadt, K. Hirai, K. Roeder, K. J. Brennand, N. Katsanis, E. Domenici, B. Devlin, P. Sklar, Gene expression elucidates functional impact of polygenic risk for schizophrenia. Nature neuroscience 19, 1442–1453 (2016).

9. A. E. Jaffe, R. E. Straub, J. H. Shin, R. Tao, Y. Gao, L. Collado Torres, T. Kam-Thong, H. S. Xi, J. Quan, Q. Chen, C. Colantuoni, W. S. Ulrich, B. J. Maher, A. Deep-Soboslay, T. B. Consortium, A. Cross, N. J. Braindon, J. T. Leek, T. M. Hyde, J. E. Kleinman, D. R. Weinberger, Developmental And Genetic Regulation Of The Human Cortex Transcriptome In Schizophrenia. bioRxiv, (2017).

10. G. Pergola, P. Di Carlo, E. D’Ambrosio, B. Gelao, L. Fazio, M. Papalino, A. Monda, G. Scozia, B. Pietrangelo, M. Attrotto, J. A. Apud, Q. Chen, V. S. Mattay, A. Rampino, G. Caforio, D. R. Weinberger, G. Blasi, A. Bertolino, DRD2 co-expression network and a related polygenic index predict imaging, behavioral and clinical phenotypes linked to schizophrenia. Translational psychiatry 7, e1006 (2017).

11. S. Freytag, J. Gagnon-Bartsch, T. P. Speed, M. Bahlo, Systematic noise degrades gene co-expression signals but can be corrected. BMC bioinformatics 16, 309 (2015).

12. A. E. Jaffe, R. Tao, A. L. Norris, M. Kealhofer, A. Nellore, J. H. Shin, D. Kim, Y. Jia, T. M. Hyde, J. E. Kleinman, R. E. Straub, J. T. Leek, D. R. Weinberger, qSVA framework for RNA quality correction in differential expression analysis. Proceedings of the National Academy of Sciences of the United States of America 114, 7130–7135 (2017).

13. B. Zhang, S. Horvath, A general framework for weighted gene co-expression network analysis. Statistical applications in genetics and molecular biology 4, Article17 (2005).

14. G. Pergola, P. Di Carlo, I. Andriola, B. Gelao, S. Torretta, M. T. Attrotto, L. Fazio, A. Raio, D. Albergo, R. Masellis, A. Rampino, G. Blasi, A. Bertolino, Combined effect of genetic variants in the GluN2B coding gene (GRIN2B) on prefrontal function during working memory performance. Psychological medicine 46, 1135–1150 (2016).

15. R. Rosenheck, J. Doyle, D. Leslie, A. Fontana, Changing environments and alternative perspectives in evaluating the cost-effectiveness of new antipsychotic drugs. Schizophrenia bulletin 29, 81–93 (2003).

16. A. Bertolino, G. Caforio, G. Blasi, M. De Candia, V. Latorre, V. Petruzzella, M. Altamura, G. Nappi, S. Papa, J. H. Callicott, V. S. Mattay, A. Bellomo, T. Scarabino, D. R. Weinberger, M. Nardini, Interaction of COMT (Val(108/158)Met) genotype and olanzapine treatment on prefrontal cortical function in patients with schizophrenia. The American journal of psychiatry 161, 1798–1805 (2004).

17. E. Radulescu, A. E. Jaffe, R. E. Straub, Q. Chen, J. H. Shin, T. M. Hyde, J. E. Kleinman, D. R. Weinberger, Identification and prioritization of gene sets associated with schizophrenia risk by co-expression network analysis in human brain. bioRxiv, (2018).

18. P. Langfelder, R. Luo, M. C. Oldham, S. Horvath, Is my network module preserved and reproducible? PLoS computational biology 7, e1001057 (2011).

19. M. R. Johnson, J. Behmoaras, L. Bottolo, M. L. Krishnan, K. Pernhorst, P. L. M. Santoscoy, T. Rossetti, D. Speed, P. K. Srivastava, M. Chadeau-Hyam, N. Hajji, A. Dabrowska, M. Rotival, B. Razzaghi, S. Kovac, K. Wanisch, F. W. Grillo, A. Slaviero, S. R. Langley, K. Shkura, P. Roncon, T. De, M. Mattheisen, P. Niehusmann, T. J. O’Brien, S. Petrovski, M. von Lehe, P. Hoffmann, J. Eriksson, A. J. Coffey, S. Cichon, M. Walker, M. Simonato, B. Danis, M. Mazzuferi, P. Foerch, S. Schoch, V. De Paola, R. M. Kaminski, V. T. Cunliffe, A. J. Becker, E. Petretto, Systems genetics identifies Sestrin 3 as a regulator of a proconvulsant gene network in human epileptic hippocampus. Nature communications 6, 6031 (2015).

20. P. Michalak, Coexpression, coregulation, and cofunctionality of neighboring genes in eukaryotic genomes. Genomics 91, 243–248 (2008).

21. G. Kustatscher, P. Grabowski, J. Rappsilber, Pervasive coexpression of spatially proximal genes is buffered at the protein level. Molecular systems biology 13, 937 (2017).

22. C. A. de Leeuw, J. M. Mooij, T. Heskes, D. Posthuma, MAGMA: generalized gene-set analysis of GWAS data. PLoS computational biology 11, e1004219 (2015).

23. X. Xu, A. B. Wells, D. R. O’Brien, A. Nehorai, J. D. Dougherty, Cell type-specific expression analysis to identify putative cellular mechanisms for neurogenetic disorders. The Journal of neuroscience: the official journal of the Society for Neuroscience 34, 1420–1431 (2014).

24. K. Ohi, T. Shimada, Y. Nitta, H. Kihara, H. Okubo, T. Uehara, Y. Kawasaki, Specific gene expression patterns of 108 schizophrenia-associated loci in cortex. Schizophrenia research 174, 35–38 (2016).

25. D. R. Weinberger, From neuropathology to neurodevelopment. Lancet 346, 552–557 (1995).

26. M. R. Johnson, K. Shkura, S. R. Langley, A. Delahaye-Duriez, P. Srivastava, W. D. Hill, O. J. Rackham, G. Davies, S. E. Harris, A. Moreno-Moral, M. Rotival, D. Speed, S. Petrovski, A. Katz, C. Hayward, D. J. Porteous, B. H. Smith, S. Padmanabhan, L. J. Hocking, J. M. Starr, D. C. Liewald, A. Visconti, M. Falchi, L. Bottolo, T. Rossetti, B. Danis, M. Mazzuferi, P. Foerch, A. Grote, C. Helmstaedter, A. J. Becker, R. M. Kaminski, I. J. Deary, E. Petretto, Systems genetics identifies a convergent gene network for cognition and neurodevelopmental disease. Nature neuroscience 19, 223–232 (2016).

27. F. Zambelli, G. Pesole, G. Pavesi, Pscan: finding over-represented transcription factor binding site motifs in sequences from co-regulated or co-expressed genes. Nucleic acids research 37, W247–252 (2009).

28. A. Ultsch, J. Lotsch, What do all the (human) micro-RNAs do? BMC genomics 15, 976 (2014).

29. M. E. Hauberg, M. H. Holm-Nielsen, M. Mattheisen, A. L. Askou, J. Grove, A. D. Borglum, T. J. Corydon, Schizophrenia risk variants affecting microRNA function and site-specific regulation of NT5C2 by miR-206. European neuropsychopharmacology: the journal of the European College of Neuropsychopharmacology 26, 1522–1526 (2016).

30. Y. Kim, P. Giusti-Rodriguez, J. J. Crowley, J. Bryois, R. J. Nonneman, A. K. Ryan, C. R. Quackenbush, M. D. Iglesias-Ussel, P. H. Lee, W. Sun, F. P. de Villena, P. F. Sullivan, Comparative genomic evidence for the involvement of schizophrenia risk genes in antipsychotic effects. Molecular psychiatry 23, 708–712 (2018).

31. D. Trabzuni, M. Ryten, R. Walker, C. Smith, S. Imran, A. Ramasamy, M. E. Weale, J. Hardy, Quality control parameters on a large dataset of regionally dissected human control brains for whole genome expression studies. Journal of neurochemistry 119, 275–282 (2011).

32. A. Schroeder, O. Mueller, S. Stocker, R. Salowsky, M. Leiber, M. Gassmann, S. Lightfoot, W. Menzel, M. Granzow, T. Ragg, The RIN: an RNA integrity number for assigning integrity values to RNA measurements. BMC Mol Biol 7, 3 (2006).

33. T. S. Stroup, J. P. McEvoy, M. S. Swartz, M. J. Byerly, I. D. Glick, J. M. Canive, M. F. McGee, G. M. Simpson, M. C. Stevens, J. A. Lieberman, The National Institute of Mental Health Clinical Antipsychotic Trials of Intervention Effectiveness (CATIE) project: schizophrenia trial design and protocol development. Schizophrenia bulletin 29, 15–31 (2003).

34. K. N. Conneely, M. Boehnke, So many correlated tests, so little time! Rapid adjustment of P values for multiple correlated tests. American journal of human genetics 81, 1158–1168 (2007).

35. L. D. Ward, M. Kellis, HaploReg: a resource for exploring chromatin states, conservation, and regulatory motif alterations within sets of genetically linked variants. Nucleic acids research 40, D930–934 (2012).

36. L. D. Ward, M. Kellis, HaploReg v4: systematic mining of putative causal variants, cell types, regulators and target genes for human complex traits and disease. Nucleic acids research 44, D877–881 (2016).

37. Z. Zhang, H. Yu, S. Jiang, J. Liao, T. Lu, L. Wang, D. Zhang, W. Yue, Evidence for Association of Cell Adhesion Molecules Pathway and NLGN1 Polymorphisms with Schizophrenia in Chinese Han Population. PloS one 10, e0144719 (2015).

38. N. The, C. Pathway Analysis Subgroup of the Psychiatric Genomics, Psychiatric genome-wide association study analyses implicate neuronal, immune and histone pathways. Nature neuroscience 18, 199–209 (2015).

39. T. Nakazawa, M. Kikuchi, M. Ishikawa, H. Yamamori, K. Nagayasu, T. Matsumoto, M. Fujimoto, Y. Yasuda, M. Fujiwara, S. Okada, K. Matsumura, A. Kasai, A. Hayata-Takano, N. Shintani, S. Numata, K. Takuma, W. Akamatsu, H. Okano, A. Nakaya, H. Hashimoto, R. Hashimoto, Differential gene expression profiles in neurons generated from lymphoblastoid B-cell line-derived iPS cells from monozygotic twin cases with treatment-resistant schizophrenia and discordant responses to clozapine. Schizophr Res 181, 75–82 (2017).

40. Y. Kim, P. Giusti-Rodriguez, J. J. Crowley, J. Bryois, R. J. Nonneman, A. K. Ryan, C. R. Quackenbush, M. D. Iglesias-Ussel, P. H. Lee, W. Sun, F. P. de Villena, P. F. Sullivan, Comparative genomic evidence for the involvement of schizophrenia risk genes in antipsychotic effects. Molecular psychiatry, (2017).

41. S. Erk, A. Meyer-Lindenberg, P. Schmierer, S. Mohnke, O. Grimm, M. Garbusow, L. Haddad, L. Poehland, T. W. Muhleisen, S. H. Witt, H. Tost, P. Kirsch, N. Romanczuk-Seiferth, B. H. Schott, S. Cichon, M. M. Nothen, M. Rietschel, A. Heinz, H. Walter, Hippocampal and frontolimbic function as intermediate phenotype for psychosis: evidence from healthy relatives and a common risk variant in CACNA1C. Biological psychiatry 76, 466–475 (2014).

42. A. Devor, O. A. Andreassen, Y. Wang, T. Maki-Marttunen, O. B. Smeland, C. C. Fan, A. J. Schork, D. Holland, W. K. Thompson, A. Witoelar, C. H. Chen, R. S. Desikan, L. K. McEvoy, S. Djurovic, P. Greengard, P. Svenningsson, G. T. Einevoll, A. M. Dale, Genetic evidence for role of integration of fast and slow neurotransmission in schizophrenia. Molecular psychiatry 22, 792–801 (2017).

43. Q. Zhang, Q. Shen, Z. Xu, M. Chen, L. Cheng, J. Zhai, H. Gu, X. Bao, X. Chen, K. Wang, X. Deng, F. Ji, C. Liu, J. Li, Q. Dong, C. Chen, The effects of CACNA1C gene polymorphism on spatial working memory in both healthy controls and patients with schizophrenia or bipolar disorder. Neuropsychopharmacology: official publication of the American College of Neuropsychopharmacology 37, 677–684 (2012).

44. B. Dietsche, H. Backes, D. Laneri, T. Weikert, S. H. Witt, M. Rietschel, J. Sommer, T. Kircher, A. Krug, The impact of a CACNA1C gene polymorphism on learning and hippocampal formation in healthy individuals: a diffusion tensor imaging study. NeuroImage 89, 256–261 (2014).

45. Psychiatric genome-wide association study analyses implicate neuronal, immune and histone pathways. Nature neuroscience 18, 199–209 (2015).

46. G. Lippi, C. C. Fernandes, L. A. Ewell, D. John, B. Romoli, G. Curia, S. R. Taylor, E. P. Frady, A. B. Jensen, J. C. Liu, M. M. Chaabane, C. Belal, J. L. Nathanson, M. Zoli, J. K. Leutgeb, G. Biagini, G. W. Yeo, D. K. Berg, MicroRNA-101 Regulates Multiple Developmental Programs to Constrain Excitation in Adult Neural Networks. Neuron 92, 1337–1351 (2016).

47. A. Jauhari, T. Singh, A. Pandey, P. Singh, N. Singh, A. K. Srivastava, A. B. Pant, D. Parmar, S. Yadav, Differentiation Induces Dramatic Changes in miRNA Profile, Where Loss of Dicer Diverts Differentiating SH-SY5Y Cells Toward Senescence. Molecular neurobiology 54, 4986–4995 (2017).

48. M. C. Chiang, Y. C. Cheng, H. M. Chen, Y. J. Liang, C. H. Yen, Rosiglitazone promotes neurite outgrowth and mitochondrial function in N2A cells via PPARgamma pathway. Mitochondrion 14, 7–17 (2014).

49. H. Chang, L. Li, T. Peng, M. Grigoroiu-Serbanescu, S. E. Bergen, M. Landen, C. M. Hultman, A. J. Forstner, J. Strohmaier, J. Hecker, T. G. Schulze, B. Muller-Myhsok, A. Reif, P. B. Mitchell, N. G. Martin, S. Cichon, M. M. Nothen, S. Jamain, M. Leboyer, F. Bellivier, B. Etain, J. P. Kahn, C. Henry, M. Rietschel, G. Swedish Bipolar Study, D. S. C. Moo, X. Xiao, M. Li, Identification of a Bipolar Disorder Vulnerable Gene CHDH at 3p21.1. Molecular neurobiology, (2016).

50. W. Sun, J. Poschmann, R. Cruz-Herrera Del Rosario, N. N. Parikshak, H. S. Hajan, V. Kumar, R. Ramasamy, T. G. Belgard, B. Elanggovan, C. C. Wong, J. Mill, D. H. Geschwind, S. Prabhakar, Histone Acetylome-wide Association Study of Autism Spectrum Disorder. Cell 167, 1385–1397 e1311 (2016).

51. M. C. O’Donovan, M. J. Owen, The implications of the shared genetics of psychiatric disorders. Nature medicine 22, 1214–1219 (2016).

52. P. Celada, M. V. Puig, F. Artigas, Serotonin modulation of cortical neurons and networks. Frontiers in integrative neuroscience 7, 25 (2013).

53. R. Brisch, A. Saniotis, R. Wolf, H. Bielau, H. G. Bernstein, J. Steiner, B. Bogerts, K. Braun, Z. Jankowski, J. Kumaratilake, M. Henneberg, T. Gos, The role of dopamine in schizophrenia from a neurobiological and evolutionary perspective: old fashioned, but still in vogue. Frontiers in psychiatry 5, 47 (2014).

54. R. Birnbaum, D. R. Weinberger, Functional neuroimaging and schizophrenia: a view towards effective connectivity modeling and polygenic risk. Dialogues in clinical neuroscience 15, 279–289 (2013).

55. J. E. Kleinman, A. J. Law, B. K. Lipska, T. M. Hyde, J. K. Ellis, P. J. Harrison, D. R. Weinberger, Genetic neuropathology of schizophrenia: new approaches to an old question and new uses for postmortem human brains. Biological psychiatry 69, 140–145 (2011).

56. A. Meyer-Lindenberg, D. R. Weinberger, Intermediate phenotypes and genetic mechanisms of psychiatric disorders. Nature reviews. Neuroscience 7, 818–827 (2006).

57. T. R. Insel, The NIMH Research Domain Criteria (RDoC) Project: precision medicine for psychiatry. The American journal of psychiatry 171, 395–397 (2014).

58. T. Moberget, N. T. Doan, D. Alnaes, T. Kaufmann, A. Cordova-Palomera, T. V. Lagerberg, J. Diedrichsen, E. Schwarz, M. Zink, S. Eisenacher, P. Kirsch, E. G. Jonsson, H. Fatouros-Bergman, L. Flyckt, KaSp, G. Pergola, T. Quarto, A. Bertolino, D. Barch, A. Meyer-Lindenberg, I. Agartz, O. A. Andreassen, L. T. Westlye, Cerebellar volume and cerebellocerebral structural covariance in schizophrenia: a multisite mega-analysis of 983 patients and 1349 healthy controls. Molecular psychiatry, (2017).

59. G. T. Consortium, Human genomics. The Genotype-Tissue Expression (GTEx) pilot analysis: multitissue gene regulation in humans. Science 348, 648–660 (2015).

60. S. R. Kay, A. Fiszbein, L. A. Opler, The positive and negative syndrome scale (PANSS) for schizophrenia. Schizophrenia bulletin 13, 261–276 (1987).

61. L. Jacob, J. A. Gagnon-Bartsch, T. P. Speed, Correcting gene expression data when neither the unwanted variation nor the factor of interest are observed. Biostatistics 17, 16–28 (2016).

62. J. A. Gagnon-Bartsch, T. P. Speed, Using control genes to correct for unwanted variation in microarray data. Biostatistics 13, 539–552 (2012).

63. D. Risso, J. Ngai, T. P. Speed, S. Dudoit, Normalization of RNA-seq data using factor analysis of control genes or samples. Nature biotechnology 32, 896–902 (2014).

64. P. Langfelder, S. Horvath, WGCNA: an R package for weighted correlation network analysis. BMC bioinformatics 9, 559 (2008).

65. P. Roussos, B. Guennewig, D. C. Kaczorowski, G. Barry, K. J. Brennand, Activity-Dependent Changes in Gene Expression in Schizophrenia Human-Induced Pluripotent Stem Cell Neurons. JAMA psychiatry 73, 1180–1188 (2016).

66. E. Ravasz, A. L. Somera, D. A. Mongru, Z. N. Oltvai, A. L. Barabasi, Hierarchical organization of modularity in metabolic networks. Science 297, 1551–1555 (2002).

67. L. J. Carithers, K. Ardlie, M. Barcus, P. A. Branton, A. Britton, S. A. Buia, C. C. Compton, D. S. DeLuca, J. Peter-Demchok, E. T. Gelfand, P. Guan, G. E. Korzeniewski, N. C. Lockhart, C. A. Rabiner, A. K. Rao, K. L. Robinson, N. V. Roche, S. J. Sawyer, A. V. Segre, C. E. Shive, A. M. Smith, L. H. Sobin, A. H. Undale, K. M. Valentino, J. Vaught, T. R. Young, H. M. Moore, A Novel Approach to High-Quality Postmortem Tissue Procurement: The GTEx Project. Biopreservation and biobanking 13, 311–319 (2015).

68. D. Smedley, S. Haider, S. Durinck, L. Pandini, P. Provero, J. Allen, O. Arnaiz, M. H. Awedh, R. Baldock, G. Barbiera, P. Bardou, T. Beck, A. Blake, M. Bonierbale, A. J. Brookes, G. Bucci, I. Buetti, S. Burge, C. Cabau, J. W. Carlson, C. Chelala, C. Chrysostomou, D. Cittaro, O. Collin, R. Cordova, R. J. Cutts, E. Dassi, A. Di Genova, A. Djari, A. Esposito, H. Estrella, E. Eyras, J. Fernandez-Banet, S. Forbes, R. C. Free, T. Fujisawa, E. Gadaleta, J. M. Garcia-Manteiga, D. Goodstein, K. Gray, J. A. Guerra-Assuncao, B. Haggarty, D. J. Han, B. W. Han, T. Harris, J. Harshbarger, R. K. Hastings, R. D. Hayes, C. Hoede, S. Hu, Z. L. Hu, L. Hutchins, Z. Kan, H. Kawaji, A. Keliet, A. Kerhornou, S. Kim, R. Kinsella, C. Klopp, L. Kong, D. Lawson, D. Lazarevic, J. H. Lee, T. Letellier, C. Y. Li, P. Lio, C. J. Liu, J. Luo, A. Maass, J. Mariette, T. Maurel, S. Merella, A. M. Mohamed, F. Moreews, I. Nabihoudine, N. Ndegwa, C. Noirot, C. Perez-Llamas, M. Primig, A. Quattrone, H. Quesneville, D. Rambaldi, J. Reecy, M. Riba, S. Rosanoff, A. A. Saddiq, E. Salas, O. Sallou, R. Shepherd, R. Simon, L. Sperling, W. Spooner, D. M. Staines, D. Steinbach, K. Stone, E. Stupka, J. W. Teague, A. Z. Dayem Ullah, J. Wang, D. Ware, M. Wong-Erasmus, K. Youens-Clark, A. Zadissa, S. J. Zhang, A. Kasprzyk, The BioMart community portal: an innovative alternative to large, centralized data repositories. Nucleic acids research 43, W589–598 (2015).

69. A. Mathelier, O. Fornes, D. J. Arenillas, C. Y. Chen, G. Denay, J. Lee, W. Shi, C. Shyr, G. Tan, R. Worsley-Hunt, A. W. Zhang, F. Parcy, B. Lenhard, A. Sandelin, W. W. Wasserman, JASPAR 2016: a major expansion and update of the open-access database of transcription factor binding profiles. Nucleic acids research 44, D110–115 (2016).

70. V. Agarwal, G. W. Bell, J. W. Nam, D. P. Bartel, Predicting effective microRNA target sites in mammalian mRNAs. eLife 4, (2015).

71. N. Wong, X. Wang, miRDB: an online resource for microRNA target prediction and functional annotations. Nucleic acids research 43, D146–152 (2015).

72. S. Bandyopadhyay, R. Mitra, TargetMiner: microRNA target prediction with systematic identification of tissue-specific negative examples. Bioinformatics 25, 2625–2631 (2009).

73. I. S. Vlachos, M. D. Paraskevopoulou, D. Karagkouni, G. Georgakilas, T. Vergoulis, I. Kanellos, I. L. Anastasopoulos, S. Maniou, K. Karathanou, D. Kalfakakou, A. Fevgas, T. Dalamagas, A. G. Hatzigeorgiou, DIANA-TarBase v7.0: indexing more than half a million experimentally supported miRNA:mRNA interactions. Nucleic acids research 43, D153–159 (2015).

74. A. L. Price, A. Helgason, G. Thorleifsson, S. A. McCarroll, A. Kong, K. Stefansson, Single-tissue and crosstissue heritability of gene expression via identity-by-descent in related or unrelated individuals. PLoS genetics 7, e1001317 (2011).

75. R. B. Scharpf, R. A. Irizarry, M. E. Ritchie, B. Carvalho, I. Ruczinski, Using the R Package crlmm for Genotyping and Copy Number Estimation. Journal of statistical software 40, 1–32 (2011).

76. B. N. Howie, P. Donnelly, J. Marchini, A flexible and accurate genotype imputation method for the next generation of genome-wide association studies. PLoS genetics 5, e1000529 (2009).

77. O. Delaneau, C. Coulonges, J. F. Zagury, Shape-IT: new rapid and accurate algorithm for haplotype inference. BMC bioinformatics 9, 540 (2008).

78. S. Purcell, B. Neale, K. Todd-Brown, L. Thomas, M. A. Ferreira, D. Bender, J. Maller, P. Sklar, P. I. de Bakker, M. J. Daly, P. C. Sham, PLINK: a tool set for whole-genome association and population-based linkage analyses. American journal of human genetics 81, 559–575 (2007).

79. J. O’Connell, D. Gurdasani, O. Delaneau, N. Pirastu, S. Ulivi, M. Cocca, M. Traglia, J. Huang, J. E. Huffman, I. Rudan, R. McQuillan, R. M. Fraser, H. Campbell, O. Polasek, G. Asiki, K. Ekoru, C. Hayward, A. F. Wright, V. Vitart, P. Navarro, J. F. Zagury, J. F. Wilson, D. Toniolo, P. Gasparini, N. Soranzo, M. S. Sandhu, J. Marchini, A general approach for haplotype phasing across the full spectrum of relatedness. PLoS genetics 10, e1004234 (2014).

80. B. Howie, C. Fuchsberger, M. Stephens, J. Marchini, G. R. Abecasis, Fast and accurate genotype imputation in genome-wide association studies through pre-phasing. Nature genetics 44, 955–959 (2012).

81. C. Genomes Project, G. R. Abecasis, A. Auton, L. D. Brooks, M. A. DePristo, R. M. Durbin, R. E. Handsaker, H. M. Kang, G. T. Marth, G. A. McVean, An integrated map of genetic variation from 1,092 human genomes. Nature 491, 56–65 (2012).

82. B. Howie, J. Marchini, M. Stephens, Genotype imputation with thousands of genomes. G3 1, 457–470 (2011).

83. O. Delaneau, J. Marchini, J. F. Zagury, A linear complexity phasing method for thousands of genomes. Nature methods 9, 179–181 (2012).

84. D. Kim, G. Pertea, C. Trapnell, H. Pimentel, R. Kelley, S. L. Salzberg, TopHat2: accurate alignment of transcriptomes in the presence of insertions, deletions and gene fusions. Genome biology 14, R36 (2013).

85. Y. Liao, G. K. Smyth, W. Shi, featureCounts: an efficient general purpose program for assigning sequence reads to genomic features. Bioinformatics 30, 923–930 (2014).

86. E. Eisenberg, E. Y. Levanon, Human housekeeping genes, revisited. Trends in genetics: TIG 29, 569–574 (2013).

87. L. Peixoto, D. Risso, S. G. Poplawski, M. E. Wimmer, T. P. Speed, M. A. Wood, T. Abel, How data analysis affects power, reproducibility and biological insight of RNA-seq studies in complex datasets. Nucleic acids research 43, 7664–7674 (2015).

88. Network, C. Pathway Analysis Subgroup of the Psychiatric Genomics, Corrigendum: Psychiatric genome-wide association study analyses implicate neuronal, immune and histone pathways. Nature neuroscience 18, 1861 (2015).

